# Mutations in the IgG B cell receptor associated with class-switched B cell lymphomas

**DOI:** 10.1101/2024.04.12.585865

**Authors:** Laabiah Wasim, Sin Wah Tooki Chu, Ben Sale, Lucy Pickard, Simon Léonard, Lingling Zhang, Helena Tolarová, Zhang Sung Tean, Niklas Engels, Dinis P. Calado, Karin Tarte, Jessica Okosun, Francesco Forconi, Pavel Tolar

## Abstract

Immunoglobulin class-switching from IgM to IgG enhances B cell receptor (BCR) signalling^1,2^ and promotes germinal centre (GC) B cell responses to antigens^3,4^. In contrast, non-Hodgkin lymphomas derived from GC B cells typically avoid IgG BCR expression and retain the unswitched IgM BCR, suggesting that the IgG BCR may protect B cells from malignant transformation^5,6^. However, the mechanism of this phenomenon and its significance for the pathogenicity of IgG-expressing lymphomas remains unclear. Here, we report that IgG-positive follicular lymphoma (FL) and the related EZB subset of diffuse large B cell lymphoma (DLBCL) acquire mutations in the IgG heavy chain, disrupting its unique intracellular tail. Enforced class switching of IgM-expressing EZB DLBCL cell lines to IgG reduces BCR surface levels, signalling via phosphoinositide-3 kinase (PI3K), levels of MYC, cell proliferation and in vivo growth. Inhibiting GSK3, a target of BCR-PI3K signalling, or stimulating the BCR rescues IgG^+^ cell proliferation. In contrast, IgG tail-truncating mutations enhance BCR surface expression, intracellular signalling and competitive growth. These findings suggest that the expansion of IgG-switched GC-like B lymphoma cells is limited by low tonic PI3K activity of the wild-type IgG BCR, but a subset of these cancers acquires mutations of the IgG intracellular tail that reverse this effect, promoting the oncogenicity of their BCRs. The presence of IgG tail mutations underscores the importance of isotype-specific BCR signalling in the pathogenesis of FL and EZB DLBCL and can potentially inform therapeutic targeting with BCR signalling inhibitors or antibody-drug conjugates.

## Introduction

FL and DLBCL are common haematological malignancies derived from GC B cells, and their pathogenesis is strongly linked to BCR activity^5,6^. Although chemotherapy combined with pan-B cell depletion is generally effective against these cancers, up to 40% of patients develop a chemotherapy-refractory or relapsing disease with poor prognosis^7^. Targeting intracellular BCR signalling pathways by small-molecule drugs or extracellular BCR domains by antibody-drug conjugates has offered new avenues for treatment. However, the effective deployment of these therapies is limited by the heterogeneity of the disease, which encompasses diverse BCR expression, BCR signalling mechanisms and drug resistance^7–10^.

One poorly understood heterogeneity in FL and DLBCL is the BCR immunoglobulin (Ig) isotype generated by class-switch recombination. Although physiological GC selection is dominated by IgG^+^ B cells (making up to 80% of antigen-specific clones)^11^, B lymphomas typically depend on the expression of the unswitched IgM BCR, with only a proportion expressing class-switched BCRs of the IgG or IgA isotypes. The IgM preference is most prominent in the activated B-cell (ABC) type DLBCL subsets MCD, A53, BN2 and N1, which express IgM in >80% of cases^5,7,8,12^. These ABC subsets are also the most aggressive and sensitive to BCR-targeted therapies^13^. Preference for IgM over normal GC B cells is, however, also apparent in FL (∼60 % of cases)^6,14,15^ and to a lower degree in the related GC B cell (GCB) type DLBCL subset EZB (∼50% of cases)^8,12^. In contrast, the GCB subset ST2 predominantly expresses IgG and IgA BCRs^8^, illustrating that the usage of the BCR isotype depends on the oncogenic background. The evasion of class-switching in FL and ABC DLBCL is underlined by frequent mutations in the Sμ switch regions that maintain IgM BCR expression despite class-switch recombination of the non-productive Ig heavy (IgH) chain allele^16,17^. This “allelic paradox” suggests that the expression of class-switched BCRs suppresses the generation of specific lymphoma subsets from class-switched B cells^6^. However, whether this suppression involves a bias in the fate choices of class-switched lymphoma-precursor B cells, for example, to differentiate into non-proliferating plasma cells (PCs) versus to re-enter a GC, or directly inhibits the growth of transformed lymphoma cells remains unclear^5,6^.

Strikingly, the preference for IgM BCR expression in specific lymphoma subsets contrasts with the positive role of the IgG BCR in enhancing the antigen-driven expansion of IgG-switched GC B cells and IgG antibody production^4,11,18^. Compared to the IgM BCR, the membrane form of the IgG heavy chain includes an extended intracellular tail, which contains the immunoglobulin tail tyrosine (ITT) motif^2^. After antigen binding, the ITT becomes phosphorylated by SYK and recruits the adaptor proteins GRB2 or GRAP to enhance signalling through intracellular Ca^2+^ and mitogen-activated protein kinase (MAPK) pathways^19,20^. Removing the tail or mutating the ITT motif abrogates the preferential production of IgG^+^ B cells, IgG^+^ PCs and IgG antibodies^4,1^. Conversely, a hyper-active ITT motif is associated with elevated auto-antibody levels and autoimmunity^21^. The IgG BCR may also promote GC B cells through its extracellular domains, although the basis for this remains unknown^11^.

Understanding the mechanisms by which the IgG BCR either promotes or inhibits lymphomagenesis compared to the IgM BCR may clarify the preference of BCR isotype usage across lymphoma subtypes and allow us to understand better the heterogeneity of oncogenic BCR signalling in the pathogenicity of the disease. Here, we show that a subset of IgG+ GC-derived lymphomas acquire mutations that truncate the IgG intracellular tail. These mutations promote the growth of IgG+ lymphoma cells by eliminating a previously unappreciated role of the IgG tail in downmodulating the surface BCR.

## Results

### Recurrent lymphoma mutations abrogate the IgG intracellular tail

To provide insights into the role of class-switching in oncogenic BCR signalling, we analysed the sequences of IgH constant regions in RNAseq and whole-exome sequencing data from four primary lymphoma datasets (EGAD00001003600^22^, NCICCR-DLBCL^12^, TCGA-DLBC (https://www.cancer.gov/tcga), MALY-DE^23^) and a collection of B lymphoma cell lines^24^. We detected recurrent missense and nonsense mutations in *IGHG1*, *IGHG3* and *IGHG4* transcripts that occurred in the last exon encoding the intracellular portions of the membrane forms of IgG1, IgG3 and IgG4, respectively (Fig. 1a, Supplementary Table 1). The mutations predominantly introduced premature stop codons truncating the intracellular tail to a sequence identical to IgM, although some also affected the critical tyrosine of the ITT motif. Overall, this suggests that these mutations modulate BCR signalling.

**Fig. 1.**
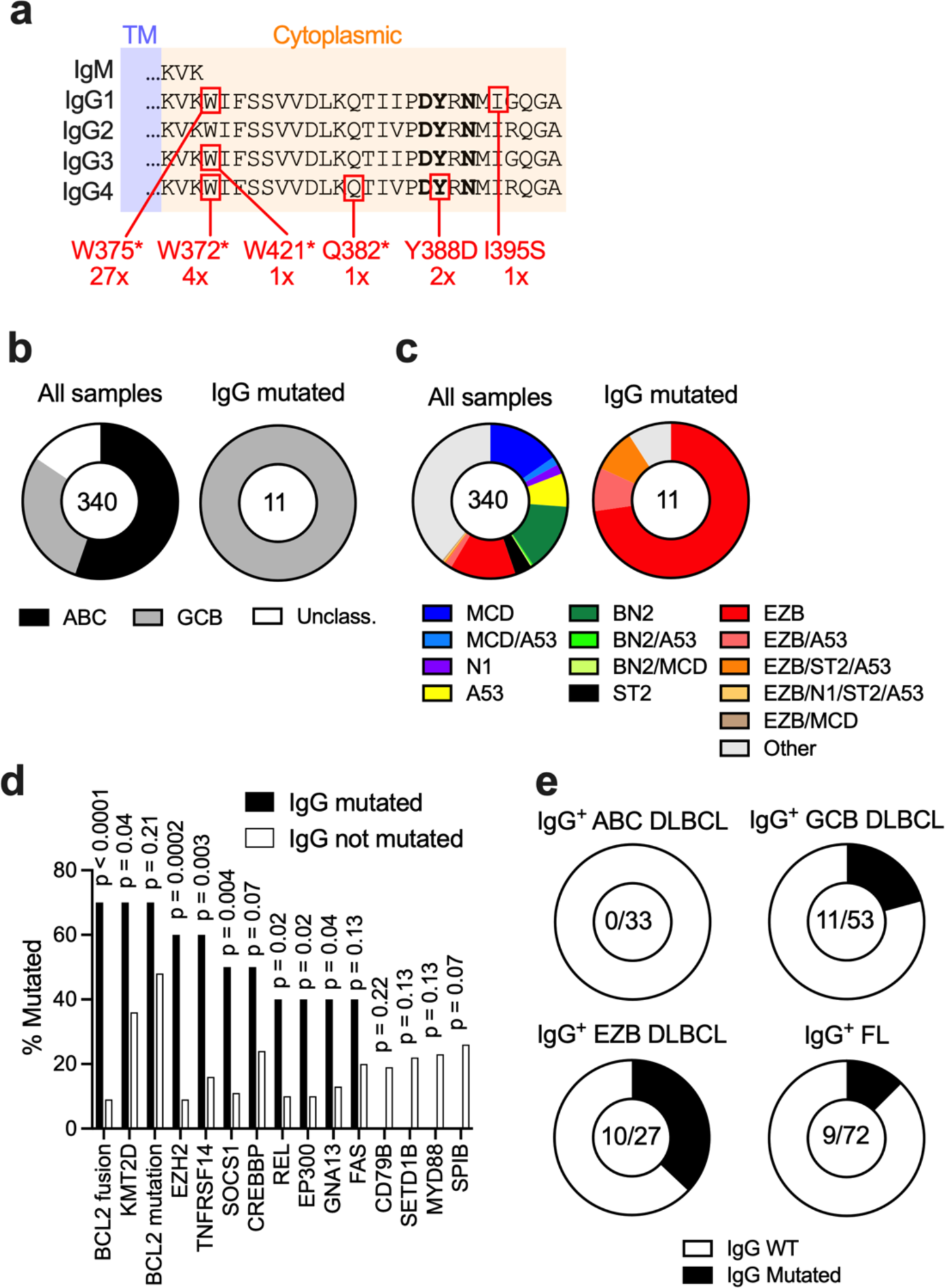
Recurrent mutations affecting the IgG cytoplasmic tail in lymphoma. (a) Summary of locations and numbers of mutations identified in all DLBCL and FL samples. The ITT motif is in bold. (b, c) Cell of origin (b) and LymphGen subset (c) classification of 340 BCR-clonal DLBCL samples in the TCGA and NCI datasets. (d) The most and least common genetic aberrations associated with IgG mutations. P, significance in Fisher’s exact test, N = 340. (e) Frequency of IgG mutations in IgG^+^ samples of the indicated subsets.

To confirm that these mutations affect the lymphoma-expressed IgG BCRs, we focused on 340 DLBCL RNAseq samples from the NCI^12^ and TCGA datasets, in which a dominant clonal IgH variable sequence could be identified and linked to the isotype of the constant region^25^. Eleven of the 104 samples expressing clonal IgG BCR (11%) contained near-clonal IgG tail mutations in the gene encoding the corresponding IgG subclass. In contrast, IgG tail mutations were absent in DLBCL samples expressing clonal IgM or IgA BCRs, a finding we validated in whole-exome sequencing data, which are not affected by RNA expression. The mutations were also absent in samples from reactive tonsils. Thus, the IgG mutations are not random events but affect the function of the cell surface BCR expressed on lymphoma cells from the productively arranged IgG heavy chain allele after class-switching.

All of the clonal IgG-mutated DLBCL samples were of the GCB subtype (Fig. 1b). LymphGen classification^8^ indicated that they are most closely associated with the EZB subset (Fig. 1c). Similarly, the most frequent genetic aberration that coincided with the IgG mutations were typical of EZB DLBCL, such as BCL2 fusion and mutations in KMT2D, EZH2 and TNFRSF14 (Fig. 1d). Within the IgG^+^ EZB samples, 37% contained the IgG tail mutations (Fig. 1e). Given the close genetic relationship of EZB DLBCL with FL and transformed FL (tFL), we also searched for IgG tail mutations in FL samples. We detected the IgG tail-truncating mutations in 3 out of 96 samples of the MALY-DE dataset, which contains whole-genome sequences corrected for germline mutations but not RNASeq to determine BCR isotype expression (Supplementary Table 1). In addition, in RNASeq samples from the FL PRIMA trial^26^, we found IgG tail mutations in 9 out of 72 (12.5%) of the IgG BCR-expressing cases, 8 of which were tail-truncating (Fig. 1e, Supplementary Table 1). Thus, a substantial fraction of IgG class-switched EZB DLBCLs and the genetically related FLs develop mutations that render the intracellular tails of their IgG BCRs similar to those of the IgM BCR.

### Class-switching to IgG inhibits the proliferation of sensitive lymphoma subsets via the intracellular tail of the membrane IgG

To understand the effect of IgM to IgG class switching on lymphoma cell growth, we used CRISPR-induced class-switching in IgM BCR-expressing cell lines representing EZB and contrasting DLBCL subsets^27^. We introduced an sgRNA targeting the IgM class-switch region (Sμ) together with an sgRNA targeting either the IgG1 (Sγ1) or IgG4 (Sγ4) switch region using a dual-sgRNA lentiviral delivery into Cas9-expressing WSU-FSCCL cells, a model of tFL/EZB DLBCL (Fig. 2a). Such dual targeting of class-switch regions resulted in approximately 10% of the cells losing their surface BCR (BCR^-^) and approximately 1.5% switching from IgM to IgG three days after transduction (Fig. 2b). The CRISPR-induced class-switching was specific because transduction with a sgRNA targeting *CD4* along with a nontargeting control sgRNA did not generate IgG^+^ cells. BCR^-^ cells were lost in subsequent cultures, confirming the BCR dependency of the cell line (Fig. 2b, c). We confirmed the BCR dependency of cell growth of the parental WSU-FSCCL cells by independent targeting of *IGHM*, which led to a gradual decrease of the targeted cells in the culture (Extended Data Fig. 2a). Importantly, WSU-FSCCL that had class-switched to IgG1 or IgG4 were also diminished in the culture over time (Fig. 2b, c). To extend the findings to other lymphoma cell lines, we induced class-switching by CRISPR in NU-DHL1 (another model of EZB DLBCL), HBL1 (a model of MCD DLBCL) and SUDHL5 (a model of ST2 DLBCL) cells (Extended Data Figure 2b-d). As seen with WSU-FSCCL cells, NU-DHL1 and HBL1 that class-switched to IgG1^+^ and IgG4^+^ were depleted from the culture over time, similarly to BCR^-^ cells (Extended Data Figure 2b, c). In contrast, IgG^+^ SUDHL5 cells remained in the culture despite the loss of BCR^-^ cells (Extended Data Figure 2d). To quantify the effects of BCR isotype on cell growth, we flow-sorted the IgG4^+^ and IgM^+^ cells for each cell line after class switching and seeded them in 1:1 ratios. The competitive co-cultures showed that IgM^+^ cells gradually outcompeted IgG4^+^ cells in WSU-FSCCL, NU-DHL1 and HBL1 cell lines, indicating that IgG class-switching is detrimental to the growth of EZB and MCD DLBCL cells (Fig. 2d). In contrast, IgG class-switched SUDHL5 cells grew with only a mild defect compared to IgM^+^ cells. To confirm the growth defect of IgG^+^ EZB DLBCL cells in vivo, we injected a ∼ 1:1 starting mixture of IgM^+^ and IgG1^+^ WSU-FSCCL or NU-DHL1 cells into NSG mice. After three weeks, IgM^+^ WSU-FSCCL cells were found in the bone marrow and IgM^+^ NU-DHL1 cells in the spleen. In contrast, only minimal numbers of IgG1^+^ cells of either cell line were recovered from these organs (Fig. 2e, Extended Data Fig. 2e). Thus, class-switching from IgM to IgG impairs the in vitro and in vivo growth of cells derived from DLBCL subsets that avoid class-switching but not those that frequently express class-switched BCRs.

**Fig. 2.**
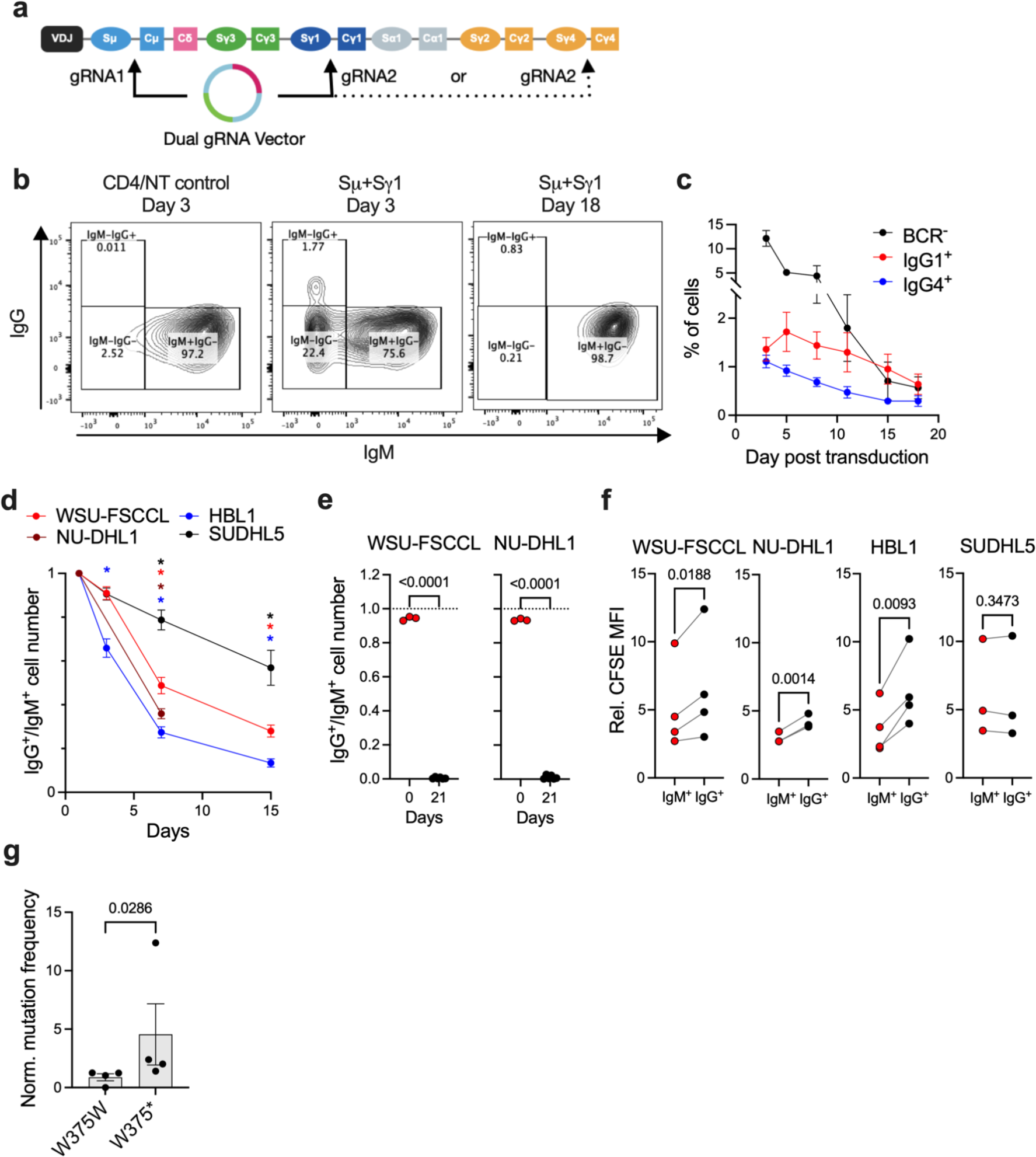
Class-switching from IgM to IgG inhibits the proliferation of EZB DLBCL cells. (a) Schematic of dual targeting of immunoglobulin switch regions to induce class-switching by CRISPR. (b) Flow-cytometry showing generation of IgM^-^ IgG^-^ (BCR^-^) and IgM^-^ IgG^+^ class-switched cells three days after CRISPR targeting of Sμ and Sγ1 compared to *CD4* and non-targeting (NT) control in WSU-FSCCL cells. Analysis at day 18 shows loss of the BCR^-^ and class-switched cells. (c) Proportions of BCR^-^ and class-switched cells over time. N = 4 experiments. (d) Relative proportions of IgG4^+^ cells in culture initially seeded at 1:1 with IgM^+^ cells. Asterisks indicate p<0.0001 in two-way ANOVA comparing proportions to day 1. N = 3 (NU-DHL1), 5 (SUDHL5) or 6 (WSU-FSCCL and HBL1). (e) Ratios of IgG1^+^ to IgM^+^ WSU-FSCCL and NU-DHL1 cells mixed approximately 1:1 before and 3 weeks after xenotransplantation into NSG mice. Ratios before transplantations are from N = 3 replicate measurements. Xenotransplant data are from the bone marrows (WSU-FSCCL) or spleens (NU-DHL1) of N = 6 mice. Brackets show significance in unpaired t-tests with Welch’s correction. (f) Dilution of CFSE after seven days of competitive culture of IgG4^+^ and IgM^+^ cells. Numbers show significance in paired t-tests. N = 3 or 4 experiments. (g) The frequency of targeted *IGHG1* alleles in IgG1^+^ WSU-FSCCL cells after five weeks of culture normalised to the initial frequency at week one. Bracket shows significance in Mann-Whitney test, N = 4 experiments. All aggregated data show means and SEM.

Antigen-stimulated normal IgG^+^ GC B cells may undergo increased Ca^2+^-induced apoptosis^28^. However, IgG class-switched WSU-FSCCL cells had similarly low levels of active caspase-3 as IgM^+^ cells (Extended Data Fig. 2f). In contrast, IgG^+^ WSU-FSCCL, NU-DHL1 and HBL1 cells had higher retention of CFSE staining after seven days of competitive culture compared to IgM^+^ cells (Fig. 2f), indicating that class-switching reduced the rate of cell division. In contrast, IgG^+^ SUDHL5 cells diluted CFSE similarly to the IgM^+^ counterparts. Thus, IgG class-switching inhibits the proliferation of cells from sensitive lymphoma subsets.

To understand if IgG tail mutations enhance the growth of IgG-switched EZB cells, we introduced the most frequent tail-truncating mutation from patient samples IgG1 W375* into the IgG1^+^ WSU-FSCCL cells using CRISPR homology-directed repair (HDR). To do so, we nucleofected cells with Cas9 and sgRNA targeting exon 6 of *IGHG1* along with an ssDNA HDR substrate that contained either a silent mutation of the PAM or the silent mutation of the PAM together with a nucleotide base change leading to the W375* mutation. We then monitored changes in the frequency of the mutations from one week to five weeks post-editing by amplicon sequencing of genomic DNA. Although it can be expected that only half of the induced mutations occur on the productive IgH allele, we observed that the frequency of the W375* mutation increased in the culture compared to the frequency of the control edits (Fig. 2f). Thus, the elimination of the IgG tail by the patient-derived mutation enhances the competitive growth of class-switched EZB lymphoma cells, consistent with the idea that these mutations promote IgG^+^ lymphoma pathogenesis.

### Class-switching to IgG reduces “toncogenic” BCR signalling in EZB DLBCL

Previously, it has been hypothesised that the IgG BCR inhibits lymphoma cell proliferation through enhanced signalling, leading to terminal differentiation into PCs^5,6^. However, CRISPR targeting of the master regulator of PC fate, PRDM1, did not rescue the growth of IgG^+^ WSU-FSCCL cells, nor did it lead to an accumulation of PRDM1-deleterious indels in the culture over time (Extended data Fig. 3a). Transcriptionally, the IgG^+^ WSU-FSCCL cells remained similar to IgM^+^ cells and did not upregulate genes associated with PC differentiation, such as PRDM1, XBP1 or SDC1 (Extended Data Fig. 3b, c, Supplementary Table 2). The IgG^+^ cells had altered expression of genes regulating BCR signalling, with the BCR pathway being the most differentially expressed gene set (Extended Data Fig. 3d). For example, the IgG^+^ cells upregulated the kinases SYK and BTK and downregulated the adaptor CD19, the transcription factor NFATC1 and phosphatases PTPN6 (SHP1) and INPP5D (SHIP1) (Extended Data Fig. 3e). However, these changes were small and did not suggest an apparent enhancement or inhibition of BCR signalling.

To understand whether class-switching alters the genetic dependency of these cells, we carried out a whole-genome CRISPR screen for genes regulating cell growth of the IgG^+^ WSU-FSCCL cells and compared the results to the parental IgM^+^ WSU-FSCCL cells. Gene CRISPR scores in IgG^+^ and IgM^+^ cells generally correlated, indicating that the IgG^+^ cells retain gene dependency similar to the parental IgM^+^ cells (Extended Data Fig. 3f, Supplementary Table 3). For example, both IgG^+^ and IgM^+^ WSU-FSCCL showed a strong dependency on the core BCR signalling components CD79A, CD79B, LYN and SYK, the CD19-CD81 complex, and on PIK3CD, consistent with the importance of BCR tonic (or “toncogenic”) signalling through PI3K in GCB DLBCL^29^. In contrast, regulators of “chronic active” BCR signalling, such as BTK, BLNK, PLCG2, PRKCB or CARD11, had no effect in either IgM^+^ or IgG^+^ cells, nor did regulators of PC differentiation PRDM1 and XBP1 (Extended Data Fig. 3f). Thus, class-switching from IgM to IgG does not result in suppressive BCR signalling or PC differentiation.

To directly determine how class-switching alters oncogenic BCR signalling in EZB versus other DLBCL subsets, we analysed the activity of the BCR pathway before and after BCR crosslinking with anti-Ig light chain antibodies. Contrary to expectations^2,19^, intracellular staining and flow cytometry of WSU-FSCCL cells showed that the IgG^+^ cells had significantly lower levels of phosphorylated CD79A, CD19, AKT and ERK compared to IgM^+^ cells both at the basal state and after BCR crosslinking (Fig. 3a). IgG^+^ NU-DHL1 cells had similar phosphorylation to IgM^+^ cells in the resting state but mildly reduced phosphorylation after BCR crosslinking (Extended Data Fig. 3g). In SUDHL5 cells, phosphorylation was also mildly, although insignificantly decreased in IgG^+^ cells compared to IgM^+^ irrespective of BCR crosslinking (Extended Data Fig. 3h). In contrast, IgG^+^ HBL1 cells had similar phosphorylation compared to IgM^+^ cells in resting state, but increased phosphorylation after BCR crosslinking (Extended Data Fig. 3i). Consistently with the phosphorylation, the BCR-induced Ca^2+^ influx was reduced in IgG^+^ WSU-FSCCL and NU-DHL1 cells compared to IgM^+^ cells (Fig. 3b and Extended Data Fig. 3j), whereas it was similar between IgM^+^ and IgG^+^ SUDHL5 cells (Extended Data Fig. 3k), and increased in IgG^+^ HBL1 cells compared to IgM^+^ cells (Extended Data Fig. 3l). Thus, class-switching modulates BCR signalling differently in each lymphoma context, decreasing BCR signalling in EZB DLBCL cells and increasing in the MCD DLBCL cells.

**Fig. 3.**
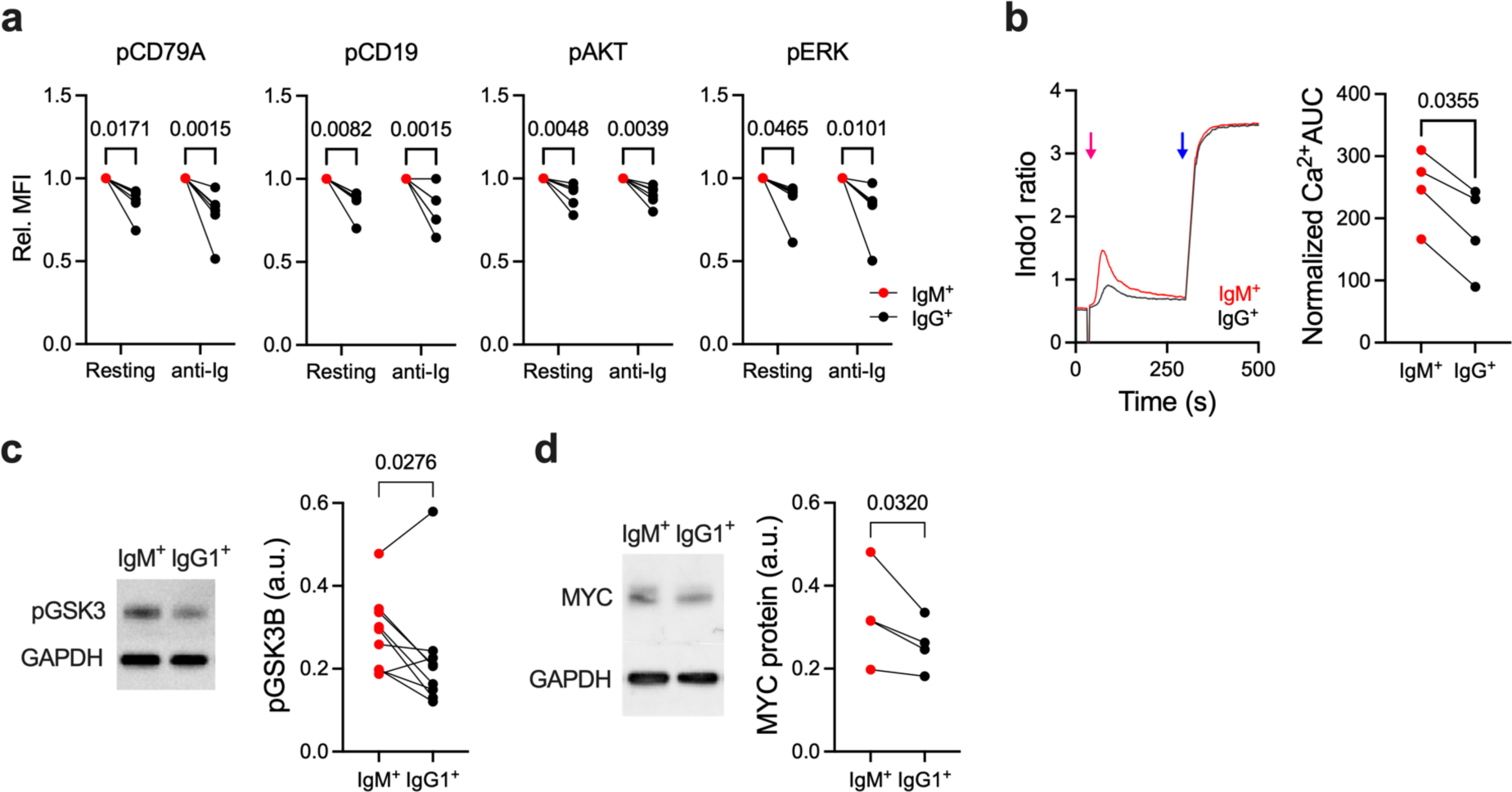
Class-switching to IgG reduces BCR oncogenic signalling. (a) Comparison of the levels of phopho-CD79A (Tyr182), phospho-CD19 (Tyr531), phospho-AKT (Ser473) and phospho-ERK1/2 (Thr202/Tyr204) in IgM^+^ and IgG4^+^ WSU-FSCCL cells before and 10 min after treatment with anti-Ig light chain antibodies as determined by intracellular staining and flow cytometry. Data show the mean fluorescence intensity of cells in each experiment normalised to values of the IgM^+^ cells. Brackets show P values from repeat-measure two-way ANOVA comparisons, N = 6 experiments. (b) Left, intracellular Ca^2+^ levels in IgM^+^ and IgG4^+^ WSU-FSCCL cells as determined by ratiometric Indo1 fluorescence before and after stimulation with anti-Ig (red arrow) and ionomycin (blue arrow). Right, quantification of the area under the Ca^2+^ curve covering anti-Ig stimulation. Brackets show P values in a paired t-test, N = 4 experiments. (c) Left, western blot detecting phosphorylated GSK3 in IgM^+^ and IgG1**^+^** WSU-FSCCL cells. Right, quantification of phospho-GSK3 (Ser9) normalised to GAPDH. P value in paired t-test, N = 9 experiments. (d) Left, western blot showing MYC protein levels. Right, quantification of MYC levels normalised to GAPDH. P value in paired t-test, N = 4 experiments.

These results suggest that reduced BCR signalling, rather than enhanced “toxic” signalling, may underlie the proliferative defect in IgG-switched EZB DLBCL cells. The proliferation of EZB DLBCL is driven by the “toncogenic” BCR-PI3K-AKT pathway, which results in the phosphorylation of an inhibitory site on GSK3 and thus inhibits GSK3-mediated phosphorylation and degradation of MYC^30^. In accord, we observed reduced GSK3 phosphorylation and reduced MYC protein levels in resting IgG^+^ WSU-FSCCL cells compared to IgM^+^ cells (Fig. 3c, d).

Importantly, bypassing the reduced tonic BCR signalling by pharmacological inhibition of GSK3 by CHIR99021 rescued the slower growth of IgG1^+^ and IgG4^+^ WSU-FSCCL and NU-DHL1 cells compared to IgM^+^ cells (Fig. 4a, Extended Data Fig. 4a-c). Moreover, CHIR99021 increased CFSE dilution in the IgG^+^ cells but not in IgM^+^ cells, indicating that GSK3 inhibition restored IgG^+^ cell proliferation (Fig. 4b, Extended Data Fig. 4a-c). In contrast, GSK3 inhibition had only a minimal effect on the growth and proliferation of IgG1^+^ HBL1 cells (Extended Data Fig. 4d). Thus, class-switching to IgG in EZB DLBCL limits proliferation by lowering BCR signalling through the PI3K-AKT-GSK3 pathway. In contrast, MCD DLBCL is affected by class-switching through a different mechanism.

**Fig. 4.**
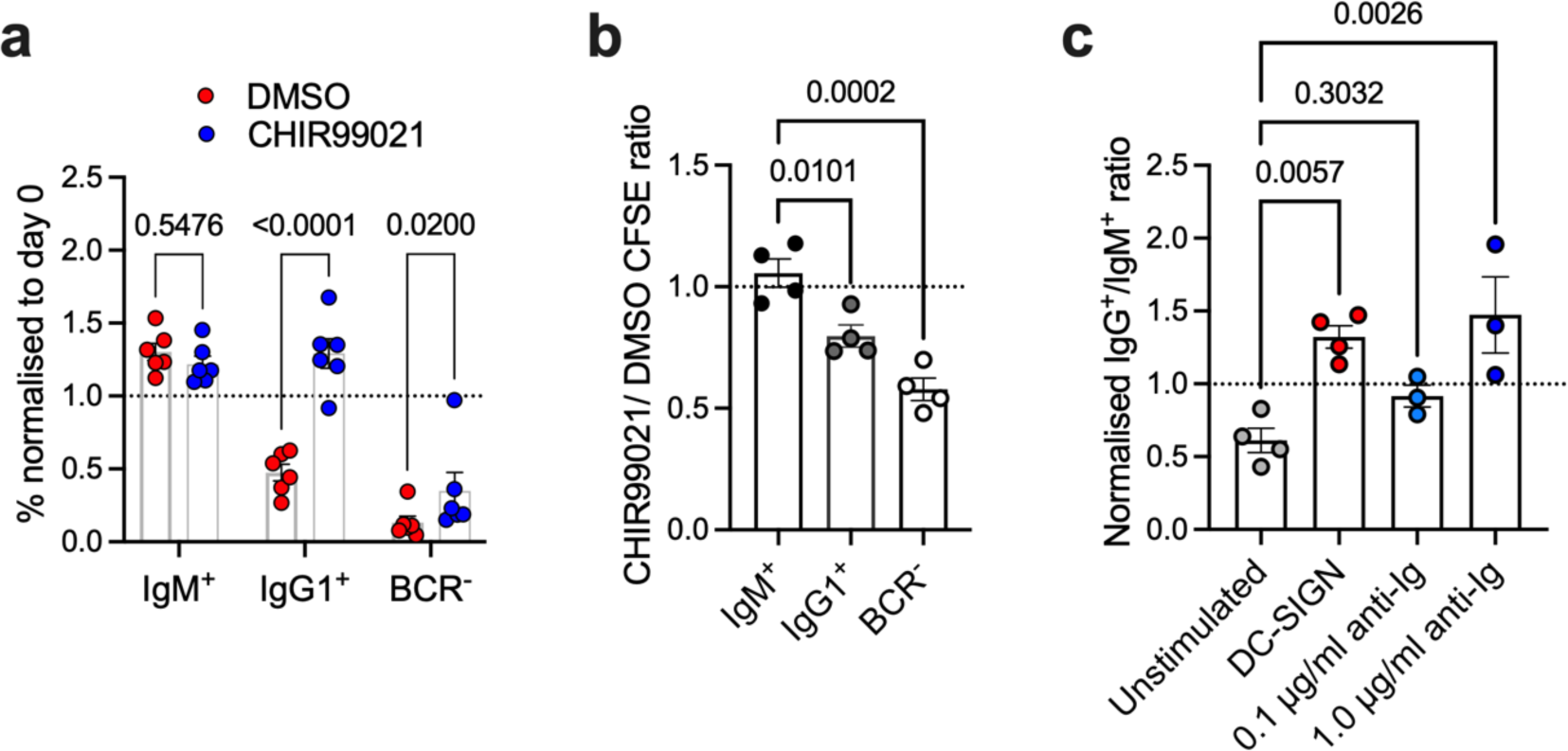
Bypassing BCR signalling to GSK3 or BCR stimulation rescues IgG-switched EZB DLBCL cells. (a) Percentage normalised to day 0 of IgM^+^, IgG1^+^ and BCR^-^ WSU-FSCCL cells 7 days after CRISPR-mediated class-switching in the presence of vehicle (DMSO) or GSK3 inhibitor (CHIR99021). Brackets show P values in two-way ANOVA, N = 6 experiments. (b) The effect of GSK3 inhibition by CHIR99021 on CFSE dilution. Brackets show P values in two-way ANOVA, N = 4 experiments. (c) Growth of IgM^+^ and IgG4^+^ WSU-FSCCL cells after stimulation. Data show the ratio of IgG^+^ to IgM^+^ cells after 7 days of stimulation, normalised to day 0. Brackets show P values in two-way ANOVA pairwise comparisons to unstimulated cells, N = 3 to 4 experiments.

To test if the effect of the BCR isotype on the growth of EZB DLBCL cells can be altered by BCR stimulation, we cultured the IgG^+^ and IgM^+^ WSU-FSCCL or NU-DHL1 cells in the absence or presence of BCR-activating ligands. Although FL and EZB DLBCL are not known to be driven by binding to cognate antigens, somatic hypermutation of the Ig variable regions in these lymphomas often introduces high-mannose glycosylation sites^25,31^, which have been proposed to stimulate growth or survival by interactions with lectins present in the tumour microenvironment. A prominent candidate for such antigen-independent BCR stimulation is DC-SIGN, a C-type lectin expressed on macrophages and FDCs in the tumour lymph nodes^25,32,33^. Both WSU-FSCCL and NU-DHL1 cells contain high-mannose glycans on their BCR and respond to DC-SIGN binding by BCR signalling^25^. To test the effect of BCR stimulation, we stimulated the IgG^+^ and IgM^+^ cells with DC-SIGN or anti-Ig in culture for seven days. The stimulation with DC-SIGN enhanced the growth of IgG^+^ WSU-FSCCL cells compared to IgM^+^ cells (Fig. 4c) and a similar, although weaker trend was observed in the NU-DHL1 cells (Extended Data Fig. 4e). In addition, the IgG^+^ cells of both cell lines were rescued with moderate 1 μg/ml doses of anti-Ig (Fig. 4c, Extended Data Fig. 4e). Thus, BCR stimulation promotes, rather than inhibits the growth of IgG^+^ EZB DLBCL cells compared to IgM^+^ cells. This result argues against the possibility that the IgG BCR generates toxic signals to EZB DLBCL lymphoma cells after ligand-induced stimulation and is consistent with the IgG^+^ cells being defective in oncogenic signalling, which is increased by DC-SIGN binding or BCR crosslinking.

### IgG tail truncation prevents BCR downmodulation and enhances PI3K signalling in EZB DLBCL

To investigate how the IgG BCR reduces oncogenic signalling in EZB lymphoma despite the presence of the signal-enhancing intracellular tail, we measured BCR surface expression. Staining for surface BCR using anti-light chain antibodies showed that IgG^+^ WSU-FSCCL, NU-DHL1 and SUDHL5 cells expressed less surface BCR than their IgM^+^ counterparts (Fig. 5a, Extended Data Fig. 5a). In contrast, BCR levels were elevated in IgG^+^ HBL1 cells compared to IgM^+^ (Extended Data Fig. 5a). This difference in surface levels was mainly due to a variation in BCR cellular distribution because total light chain staining in permeabilised IgG^+^ cells was similar to that in IgM^+^ cells in all cell lines (Fig. 5b). Thus, class-switching from IgM to IgG reduces BCR surface levels in EZB and ST2 DLBCL.

**Fig. 5.**
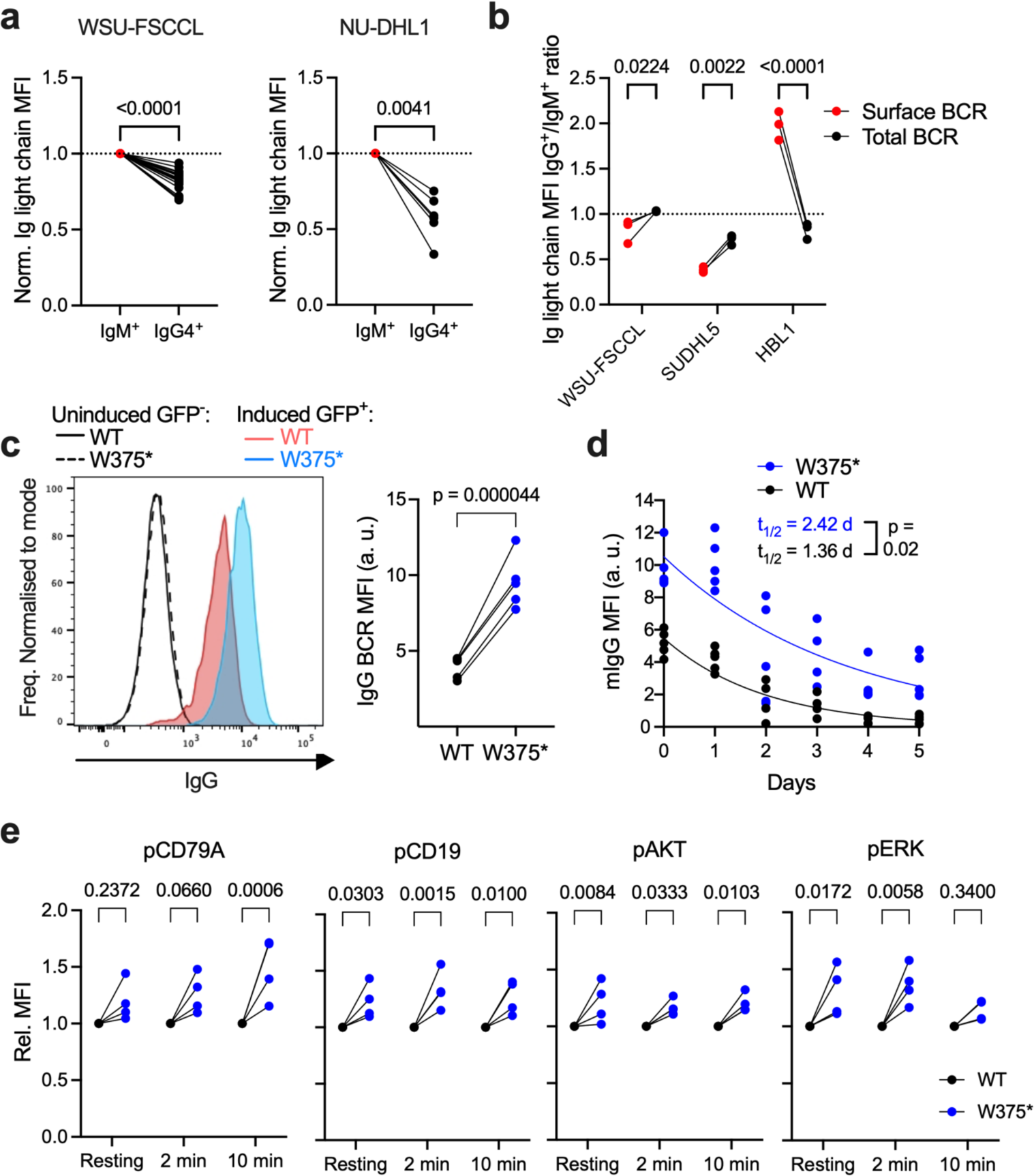
IgG tail regulates BCR surface levels and signalling. (a) Surface BCR levels in IgM^+^ and IgG4^+^ WSU-FSCCL and NU-DHL1 cells, normalised to IgM^+^ levels, as determined by anti-Ig light chain staining and flow cytometry. Brackets show P values in paired t-tests, N = 25 or 6. (b) Total Ig light chain levels in permeabilised cells compared to surface levels as determined by flow cytometry. Brackets show P values in repeat-measure two-way ANOVA comparisons, N = 3 experiments. (c) Left, representative flow cytometry showing surface IgG BCR levels in WSU-FSCCL cells transduced with constructs expressing WT or W375* mutated IgG1 containing the endogenous variable domain and bicistronic GFP. Black and dotted histograms show cells without doxycycline-induced expression, blue and red histograms show GFP^+^ cells 2 days after doxycycline-induced expression. Right, quantification of surface IgG BCR levels in GFP^+^ cells. Bracket shows significance in paired t-test. N = 5 experiments. (d) Loss of surface IgG after doxycycline removal. Lines show exponential fits, N = 5 experiments. P, significance in extra sum-of-squares F-test comparing the fitted half-lives. (e) Comparison of the levels of phopho-CD79A, phospho-CD19, phospho-AKT and phospho-ERK in GFP^+^ IgG1 WT and W375* cells before, 2 and 10 min after treatment with anti-Ig light chain antibodies as determined by intracellular staining and flow cytometry. Data show the mean fluorescence intensity of cells in each experiment normalised to values of the IgG1 WT cells. Brackets show P values from two-way repeat-measure ANOVA, N = 4 experiments.

To test if the reduction in IgG BCR levels depends on the intracellular tail, we cloned the IgH variable region from WSU-FSCCL cells and combined it with the WT constant region of membrane IgG1 or the IgG1 constant region in which the intracellular tail is truncated by the W375* mutation. We expressed these constructs linked via internal-ribosomal entry site (IRES) to GFP in WSU-FSCCL cells using a doxycycline-inducible system. Doxycycline induced similar levels of GFP (Extended Data Fig. 5b, c) and an equivalent downmodulation of the endogenous surface IgM in GFP^+^ cells from both constructs (Extended Data Fig. 5d), the latter likely due to competition for BCR components CD79A, CD79B or the Ig light chain. However, the W375* IgG1 mutant produced higher surface IgG1 expression levels than WT IgG1 (Fig. 5c). While WT and W375* IgG appeared on the cell surface with similar kinetics after induction, the W375* IgG1 mutant was lost from the cell surface at a slower rate than WT after doxycycline withdrawal (Fig. 5d). Thus, the tail-truncating mutation increases surface BCR expression in EZB DLBCL lymphomas by impairing downmodulation from the cell surface.

To test whether the enhanced surface expression of the W375* IgG1 translates into increased BCR signalling, we measured phosphorylation in the BCR pathway in GFP^+^ IgG^+^ cells by intracellular staining. Expression of W375* IgG1 led to significantly higher phospho-CD19, phospho-AKT and phospho-ERK, and modestly higher phospho-CD79A compared to WT IgG1 in resting cells (Fig. 5e). All of these signals were also increased after BCR crosslinking in cells expressing W375* compared to WT IgG. Thus, the truncating mutation of the IgG tail enhances surface BCR levels and intracellular signalling in EZB DLBCL.

## Discussion

The paradoxical avoidance of the expression of the IgG BCR in B cell lymphomas^5,6^ suggests a role for immunoglobulin class-switching in disease pathogenicity but has lacked a mechanistic explanation. One possibility is that avoidance of IgG class-switching arises early in pathogenesis because antigen-stimulated memory IgM^+^ B cells containing founder lymphoma mutations may preferentially re-enter GCs, accelerating the accumulation of additional oncogenic mutations. In contrast, antigen-stimulated IgG^+^ pre-lymphoma memory cells may preferentially differentiate into PCs^6,34,35^. However, these differences in memory B cell fates may also emerge from the association of IgM^+^ and IgG^+^ cells with different memory B cell subsets^36^. While these effects on pre-lymphoma cells will need to be investigated in dedicated in vivo models, our data show that class-switching from IgM to IgG has a cell-intrinsic oncostatic effect on cells derived from fully formed EZB and MCD lymphomas. This effect requires the WT IgG intracellular tail, which is eliminated by mutations in a subset of IgG^+^ FLs and EZB DLBCLs. Thus, class-switching to the WT IgG BCR can suppress the progression of IgG^+^ FL and EZB DLBCL through modulation of antigen-independent BCR oncogenic signalling. Mutations truncating the intracellular IgG tail break through this inhibition, allowing IgG-mutated lymphoma cells to outgrow.

Previous studies showed that the ITT motif found in the IgG intracellular tail boosts antigen-stimulated Ca^2+^ and MAPK signalling by recruiting the adaptors GRB2 or GRAP^19,20^. We observed this expected increase in anti-Ig-stimulated BCR signalling after class-switching to IgG in a model of MCD DLBCL, which depends on NF-κB signalling induced by self-antigen binding^37^. Given that class-switching in this MCD model also suppressed growth, ITT signalling from the IgG BCR may be toxic in this subset. In contrast, class-switching to IgG in EZB DLBCL suppressed cell growth by impairing “toncogenic” BCR-PI3K-AKT-GSK3 signalling caused by reduced BCR surface expression. Thus, in addition to enhancing signalling, the IgG intracellular tail also promotes downmodulation of the BCR from the cell surface, which can lead to a net loss of BCR signalling in specific contexts. The downmodulation may occur through trafficking into endosomal compartments for degradation^38^. Indeed, experimental IgG tail truncation or ITT tyrosine substitution to alanine increases BCR surface expression in mouse GC B cells^11^ and in human Burkitt lymphoma cell lines^19^, respectively. Notably, IgG^+^ FLs express lower BCR levels than normal GC B cells, which is the opposite of IgM^+^ FLs^39^. In contrast, experimental mutation of the ITT tyrosine to phenylalanine, which blocks GRB2 binding and abolishes ITT signalling, slightly reduces surface BCR levels^19^. Thus, although ITT signalling may contribute to lymphoma inhibition in certain situations, our data indicate that the IgG tail-truncating mutations in EZB DLBCL promote pro-tumour BCR activity not just by removing ITT signalling, but also by enhancing BCR surface expression. This is consistent with findings showing that “toncogenic” PI3K signalling in GCB lymphomas is highly dependent on BCR surface levels irrespectively of BCR isotype^29^.

The regulation of BCR surface levels upon class-switching may also be relevant to normal B cell responses to antigens. Although the IgG tail enhances B cell expansion and PC differentiation through the ITT motif^4,19^, it is not yet clear how exactly ITT signalling alters transcriptional wiring to contribute to this effect^40,41^. Enhanced trafficking of the IgG BCR to lysosomes^38^ may promote antigen processing and presentation on MHC II, which in turn may bestow IgG^+^ cells with higher T-follicular helper (T_fh_)-derived signals, such as CD40L and IL-21. We propose that a synergy between antigen-stimulated IgG BCR intracellular signalling and T_fh_-derived help is responsible for the enhanced expansion and differentiation of IgG^+^ cells in antigen-specific responses. For example, T_fh_-induced positive selection in the GC helps to rescue IgG^+^ GC B cells from BCR Ca^2+^-dependent apoptosis^28^. IgG^+^ B cells transforming into FL or EZB DLBCL likely lose stimulation by antigen and T_fh_ cells but remain protected from apoptosis due to BCL2 overexpression. Nevertheless, they become uncompetitive due to poor tonic or DC-SIGN-induced BCR signalling. Cells acquiring the IgG tail mutations gain a competitive advantage amongst lymphoma cells by enhancing surface BCR expression and PI3K signalling, driving accumulation of MYC and potentially other downstream effectors. Mutations mimicking T cell help could also promote the growth of WT IgG^+^ GCB lymphomas, as could be the case in the ST2 subset^8^. In contrast, ABC lymphomas use strong, self-antigen-driven BCR signalling, which may, upon IgG class-switching, trigger rapid apoptosis, preventing the acquisition of rescue mutations in the IgG tail.

Following the discovery of CD79B mutations in ABC DLBCL^42^, our study describes a second oncogenic hit to the BCR signalling domains, underscoring the importance of the BCR pathway in lymphomagenesis and corroborating its potential as a therapeutic target. The IgG mutations in FL and EZB DLBCL segregate from the CD79B mutations in ABC DLBCL, indicating that they dysregulate BCR signalling in distinct lymphoma subsets. Notably, FL and EZB DLBCL patients show heterogeneity in disease progression that is linked to the BCR. For example, patients with EZB lymphoma containing high-mannose glycosylation of the BCR show decreased progression-free survival and higher pathogenicity was suggested for IgM^+^ FL compared to IgG^+^ FL^39^. Further understanding of the role of BCR isotype and IgG mutations can help to illuminate these clinical differences and unravel the heterogeneous roles of the BCR in disease progression and response to therapy. Overall, the IgG tail mutations may signify a strong addiction of lymphoma cells to BCR signalling, which may be therapeutically exploited. Similarly, the surface expression-enhancing effect of the IgG tail mutation may sensitise these cancers to antibody-drug conjugates binding to the BCR extracellular domains, such as polatuzumab-vedotin.

## Methods

### Bioinformatic identification of IgG cytoplasmic tail mutations

RNAseq and whole genome sequencing (WGS) DLBCL datasets were obtained from the European Genome-Phenome Archive (EGAD00001003600^22^, n = 769), the NCI Genomic Data Commons Data Portal (NCICCR-DLBCL^12^, subset of n = 307 of BCR clonal samples described in^25^, and from The Cancer Genome Atlas (TCGA, https://www.cancer.gov/tcga, TCGA-DLBC, subset of n = 33 BCR clonal samples described below). RNAseq FL datasets were from the PRIMA trial^26^, n = 148. RNAseq data of B lymphoma cell lines were from EGAD00001000725^24^, n = 81, or produced in-house (HF1, WSU-FSCCL). The WGS data from n = 48 DLBCL and n = 96 FL samples and matched germlines were from the International Cancer Genome Consortium (MALY-DE^23^).

For the NCICCR-DLBCL dataset, 307 clonal IgH variable sequences and their linked constant regions were obtained as described^25^. For the TCGA-DLBCL data, variable IgH sequences and their linked constant regions were reconstructed using Trust4 (v1.0.9) and 33 samples with clonally dominant BCRs (defined as those with the most frequent IgH clone having > 5-fold higher read count over the next most abundant clone) were analysed further. For the PRIMA FL trial, IgG-expressing cases were identified using Trust4.

Initial variant calling was conducted using SAMtools mpileup (v1.17) and VarScan2 (v2.4.2). The genomic coordinates of the intracellular tail regions of all IgG isotypes were provided as a BED file and the minimum coverage was set to 5. Genomic coordinates of variants from alignments to human reference assembly GRCh37 were converted to GRCh38 using LiftOver with an Ensembl chain file. After standardisation to GRCh38, variant calls were annotated using SnpEff (v5.1d). Variants present in the Genome Aggregation Database (gnomAD v3.1.2, accessed May 2023) were filtered out. The IgG mutations were verified manually in RNASeq and, where available, exome sequencing. In some cases, VarScan2 reported mutations in two highly similar IgG subclasses (e.g. IGHG1 and IGHG3 or IGHG4 and IGHG2), but manual inspection of the aligned reads in IGV revealed that only one gene was mutated while the detection of the mutation in the other resulted from reads misaligned to a region in which the IgG sequences were identical between the subclasses. The mutations had in-sample frequencies of 60%-100% indicating that they were near-clonal (Supplementary Table 1). In the 340 DLBCL BCR clonal samples, the high mutation frequency together with the BCR clonality indicates that the mutations came from the dominant lymphoma clone. Sample TCGA-GS-A9TQ contained predominantly IgM (∼75%) but also clonally-related IgG1 (∼25%), which contained IgG1 W375* mutation. Since the mutated IgG1 subclone was subdominant, this sample was not included amongst the IgG mutated samples used in subsequent analyses. IDs of DLBCL samples with IgG WT and tail mutation were used to filter the data for analysis of associated mutations and cell-of-origin and LymphGEN^8^ subset classifications. No IgG tail mutations were detected in polyclonal B cells from 4 samples of reactive lymph nodes. The aggregated data of all the IgG mutations are shown in Fig. 1a and in Supplementary Table 1.

### Cell Lines and cell culture

WSU-FSCCL, HBL1, NU-DHL1 and SUDHL5 cell lines and their derivatives were cultured in RPMI (Gibco) supplemented with 10% heat-inactivated fetal bovine serum (FBS), 1X MEM non-essential amino acids (Gibco), 1X GlutaMAX (Gibco), 50µM 2-mercaptoethanol (Sigma Aldrich), 100U/ml penicillin and 100µg/ml streptomycin. HEK 293T cells were cultured in DMEM (Gibco) supplemented with 10% FBS, 1X MEM non-essential amino acids (Gibco), 1X GlutaMAX (Gibco), 50µM 2-mercaptoethanol (Sigma Aldrich), 100U/ml penicillin and 100µg/ml streptomycin. All cell lines were maintained at 37°C in humidified incubators with 5% CO_2_. All cell lines were authenticated and mycoplasma-tested by the Francis Crick Institute Cell Services.

### Lentiviral Production

Target CRISPR sgRNA gene sequences were designed using the Broad Institute sgRNA Designer CRISPick (Table 1). Forward and reverse oligonucleotide sequences including the guide sequence were synthesized, phosphorylated, annealed and individually cloned into vector plasmids lentiGuide-Puro (Addgene #52963) or lentiGuide-mCherry (Plasmid #170510). For CRISPR-mediated class-switching sgRNAs targetgin switch regions were cloned into a dual-targeting lentiGuide-mCherry using min U6 and min H1 promoters based on the design as described^43^. Replication-incompetent lentiviruses were produced by co-transfecting 10×10^6^ HEK293T cells with 40μg vector plasmid, 10μg envelope plasmid pMD2.G (A gift from Didier Trono (Addgene #12259)) and 30μg packaging plasmid psPAX2 (A gift from Didier Trono (Addgene #12260)) in opti-MEM media (ThermoFisher) using transIT-LT1 transfection reagent (Mirus, MIR2306). Lentivirus was harvested 48 and 72 hrs post-transfection by ultracentrifugation from cell culture supernatant (50,000 g at 4°C for 3 hours). Lentivirus production was scaled up as necessary to produce the lentiviral library.

**Table 1:**
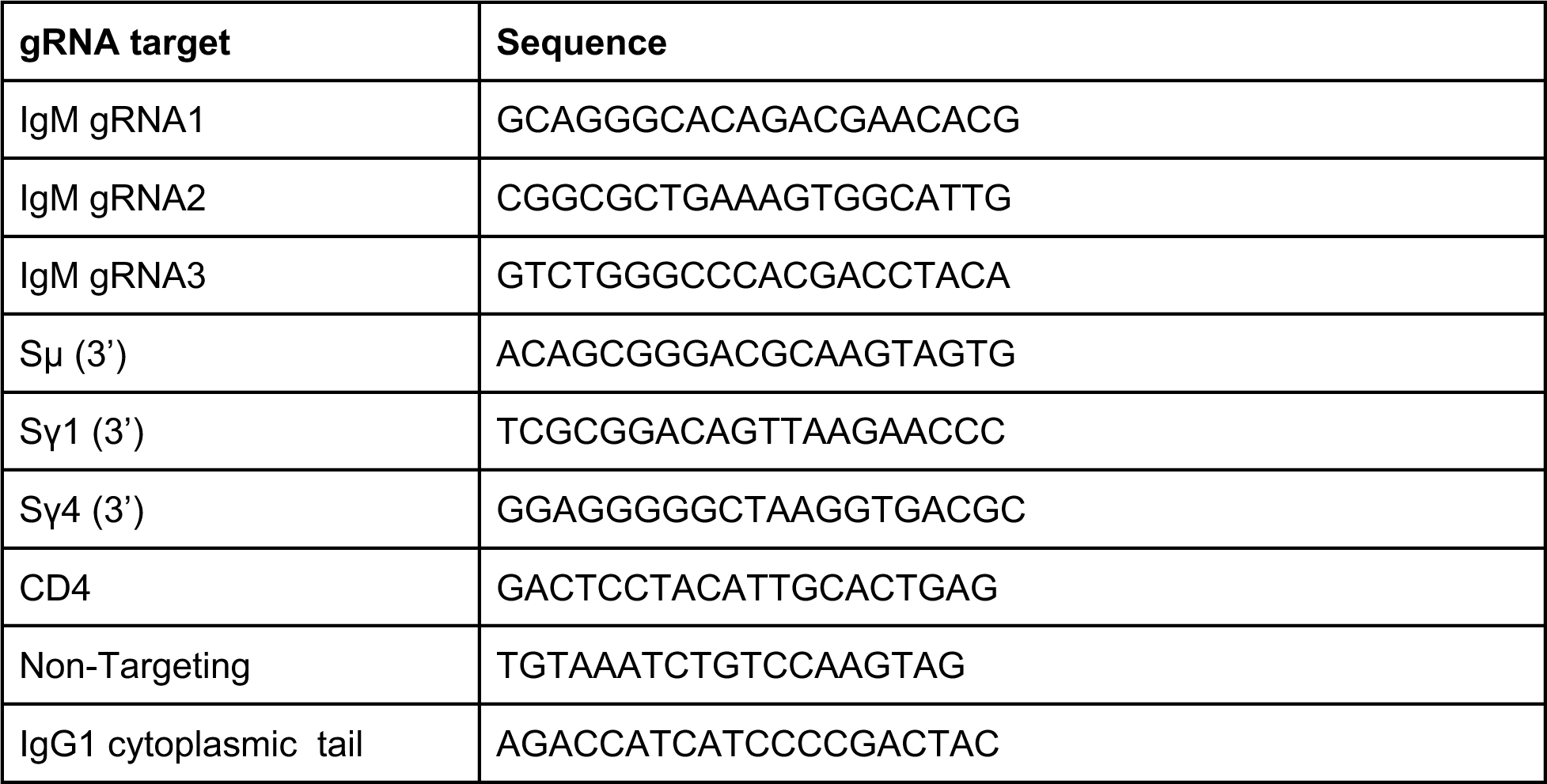
sgRNA sequences for CRISPR-mediated class-switching, knockdown, and HDR.

### Lentiviral transduction

Stable Cas9 expression was established in human B-cell lines (WSU-FSCCL, HBL-1 and SUDHL5) using lentiviral transduction of lentiCas9-Blast (Addgene #52962). Cells were incubated with 10µg/ml polybrene (Sigma-Aldrich) for 30 min before the addition of lentivirus. Cells were then spinfected at 1,300 g for 90 min at RT, following which they were incubated at 37°C. Cells were put under 10 µg/ml blasticidin selection three days post-transduction and cultured in blasticidin for 10 days, followed by single-cell sorting and confirmation of Cas9 expression by western blot.

Single-gene or dual-gene knockouts were conducted by transduction of target sequence containing lentiGuide-puro, lentiGuide-mCherry, or dual-targeting lentiGuide-mCherry virus as indicated into stable Cas9 expressing human B-cell lines by spinfection. Successfully transduced cells were then either selected in puromycin (ThermoFisher Scientific) at 1 µg/ml or detected by mCherry expression by flow cytometry.

### Ribonuclear protein nucleofection

Synthetic sgRNAs were ordered from Synthego with 2’-O-Methyl and 3’ phosphorothioate chemical modifications. Ribonuclear protein complexes were formed by incubating sgRNAs with Alt-R™ S.p. Cas9 Nuclease V3 (Integrated DNA Technologies, 1081058) at a molar ratio of 1:1 (150 pmol sgRNA with 150 pmol Cas9 enzyme per reaction) for 30 mins at 4°C. For dual targeting, sgRNA targeting Sµ (3’) and Sγ1 (3’) or Sγ4 (3’) (Table 1) were incubated together with Cas9 nuclease. NU-DHL1 cells were transfected using the SG Cell Line 4D-Nucleofector™ X Kit V4XC-3024 (Lonza) with the DS-104 setting.

### Mouse xenografts

Six weeks old male NSG mice (*NOD.Cg-Prkdc^scid^ Il2rg^tm1Wjl^/SzJ*) were tail vein-injected with 10 x 10^6^ WSU-FSCCL or 10 x 10^6^ NU-DHL1 cells each containing approximately 1:1 mixture of IgG1^+^ and IgM^+^ cells obtained by cell sorting after CRISPR-induced class-switching. Three weeks later, bone marrow and spleens were analysed for the presence of IgG^+^ and IgM^+^ WSU-FSCCL (mCherry^+^, human CD19^+^) or NU-DHL1 (human CD20^+^, human CD19^+^) cells by cell surface staining and flow cytometry (Table 2). All mice were bred and maintained at The Francis Crick Institute biological resources facility under specific pathogen-free conditions. Animal experiments were carried out in accordance with national and institutional guidelines for animal care and were approved by The Francis Crick Institute biological resources facility strategic oversight committee (incorporating the Animal Welfare and Ethical Review Body) and by the Home Office, UK.

**Table 2.** Antibodies used for immunostaining and cell stimulation. FC = Flow cytometry, S = stimulation, WB = Western Blot

| Antibody | Dilution | Use | Supplier | Catalogue Number |
| --- | --- | --- | --- | --- |
| Alexa Fluor® 647- conjugated AffiniPure Fab Fragment Goat Anti-Human IgM, Fc<sub>5</sub>μ Fragment specific | 1:500 | FC | Jackson ImmunoResearch | 109-607-043 |
| DyLight® 405- conjugated AffiniPure Fab Fragment Goat Anti-Human IgG, Fcγ Fragment specific | 1:500 | FC | Jackson ImmunoResearch | 109-477-008 |
| Anti-rabbit IgG (H+L), F(ab')<sub>2</sub> Fragment (Alexa Fluor® 488 Conjugate) | 1:1000 | FC | Cell signalling Technology | 4412 |
| Brilliant Violet 421™ anti-human CD19 Antibody | 1:200 | FC | BioLegend | 302234 |
| BV 650 anti-human CD19 Antibody | 1:50 | FC | BioLegend | 302237 |
| FITC anti-human CD20 Antibody | 1:100 |  | BioLegend | 302303 |
| Biotin Anti-Kappa light chain antibody | 1:300 | FC | Abcam | ab46807 |
| Alexa Fluor® 488 anti-human Ig light chain λ Antibody | 1:200 | FC | BioLegend | 316612 |
| Phospho-GSK-3β (Ser9) (D85E12) XP® Rabbit mAb | 1:1000 | WB | Cell signalling Technology | 5558 |
| Phospho-AKT (Ser473) Antibody | 1:50 | FC | Cell signalling Technology | 9271 |
| Phospho-p44/42 MAPK (Erk1/2) (Thr202/Tyr204) Antibody | 1:50 | FC | Cell signalling Technology | 9101 |
| Phospho-CD79A (Tyr182) Antibody | 1:50 | FC | Cell signalling Technology | 5173 |
| Phospho-CD19 (Tyr531) Antibody | 1:50 | FC | Cell signalling Technology | 3571 |
| c-Myc Antibody | 1:1000 | WB | Cell signalling Technology | 9402 |
| GAPDH (14C10) Rabbit mAb | 1:1000 | WB | Cell signalling Technology | 2118 |
| Goat F(ab')<sub>2</sub> anti Human Kappa Light Chain | As stated or 5µg/ml | S | BioRad | STAR128 |
| Goat F(ab')<sub>2</sub> anti Human Lambda Light Chain | As stated or 5µg/ml | S | BioRad | STAR130 |
| Anti-CRISPR-Cas9 antibody [7A9-3A3] | 1:1000 | WB | Abcam | ab191468 |
| IRDye® 800CW Goat anti-Rabbit IgG Secondary Antibody | 1: 10 000 | WB | LI-COR Biosciences | 926-32211 |
| IRDye® 680RD Goat anti-Mouse IgG Secondary Antibody | 1: 10 000 | WB | LI-COR Biosciences | 926-68070 |
| Goat anti-Rabbit IgG HRP conjugated | 1:10 000 | WB | Cell Signaling Technology | 7074S |

### CRISPR-mediated HDR gene editing

SgRNA targeting the IgG1 cytoplasmic tail (Table 1) was used for HDR editing in WSU-FSCCL. Short single-stranded DNA HDR templates were synthesised (Ultramer oligonucleotides, IDT) with silent mutations in PAM sites to prevent continuous editing. A double-stranded Cas9 target sequence was incorporated on the 5’ end of the IgG1 HDR templates to improve editing efficiency^44^. Cells were aliquoted into Nucleocuvette^TM^ reaction wells (20 µl/ well) and the RNP mix was added along with 2 nmol of the annealed HDR template. Cells were transfected as above using the DS-104 setting and allowed to rest at 37°C for 15 mins, then re-plated to a 48-well plate with pre-warmed media overnight before transferring to a larger volume. Cells were cultured for 5 weeks to observe the effect of the IgG cytoplasmic tail mutation on cell growth.

### Amplicon sequencing for the detection of HDR edits

Genomic DNA (gDNA) was harvested using the DNeasy Blood & Tissue Kit (Qiagen) from cells at day 7 post-transfection and week 5 post-transfection. The IgG cytoplasmic regions were PCR amplified using primers with adaptor sequences for next-generation sequencing. Forward primer IgG1 cytoplasmic tail:

5’-TCGTCGGCAGCGTCAGATGTGTATAAGAGACAGCACCAAGCCTCAGAGCAGGC - 3’

Reverse primer IgG1 cytoplasmic tail:

5’-GTCTCGTGGGCTCGGAGATGTGTATAAGAGACAGCGAGGCCAGAGAGTCATGGG - 3’

Indexed Illumina sequencing adapters were then added during a limited-cycle amplification step and pooled libraries were sequenced using an Illumina MiSeq (250 bp paired-end sequencing). The frequency of HDR editing was calculated using CRISPResso2^45^. HDR frequencies at week 5 were normalised to week 1 for each sample. Changes in frequency over time of the IgG1 W375*/PAM-mutated allele were compared to the PAM-only mutated allele.

### Whole genome CRISPR screens

Brunello sgRNA library with puromycin resistance containing 76,441 sgRNAs targeting 19,114 human genes was used to conduct the genome-wide screen in WSU-FSCCL cells. Lentiviral library virus produced in HEK293T cells was transduced by spinfection into WSU-FSCCL Cas9 cells or sorted WSU-FSCCL Cas9 IgG^+^ cells to give ∼500x library coverage at 0.3 multiplicity of infection. Cells were grown in 1 µg/ml puromycin from 72 hours post-transduction to select for successfully transduced cells. Cells were cultured at 1000x library coverage under puromycin selection for up to 4 weeks. Genomic DNA from at least 20×10^6^ cells was extracted at 3 days, 1 week, 3 weeks, and 4 weeks post-transduction using the DNeasy Blood & Tissue kit (Qiagen). Genomic DNA was also extracted from HEK293T cells used for lentiviral production to confirm baseline sgRNA frequencies. Guide RNA sequence was amplified using a two-step PCR to first amplify sgRNA sequence and then to add sequencing adaptors for Illumina sequencing. Both PCRs were performed using Phusion Flash High Fidelity Master Mix (Thermo).

PCR1 primer forward: 5’-CCCGAGGGGACCCAGAGAG-3’

PCR1 primer reverse: 5’-GCGCACCGTGGGCTTGTAC-3’

In the first PCR reaction for each sample, 10 reactions were carried out with 1.2 μg gDNA per reaction. Products from the first PCR were pooled and 10 μl amplified DNA was used in each reaction for the second PCR, carrying out 12 reactions/sample. PCR2 reverse primers contained a unique barcoded region for each sample. PCR2 products were pooled for each sample and purified using MinElute PCR Purification Kit (Qiagen). Samples were sequenced using HiSeq2500 (Illumina). Raw sequencing reads in demultiplexed FASTQ files were trimmed to contain the sgRNA sequences and mapped to the library sgRNA sequences using BowTie to calculate the frequency of each gRNA in each sample. The number of reads for each sgRNA was then normalised by the total number of reads for all gRNA in each sample to get sgRNA frequencies. Successful lentiviral library coverage across library transduction was confirmed by comparing normalised sgRNA frequencies from WSU-FSCCL samples at day 3 post-transduction to frequencies from HEK293T cells. CRISPR gene scores were calculated as:

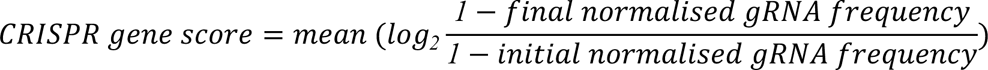

using data from day 3 and weeks 3-4. Statistics for gene level data were obtained using MAGeCK^46^, giving a ranked list of statistically significant positive and negative regulator genes. Successful targeting of genes was confirmed by comparing CRISPR gene scores and FDR values for 187 commonly essential genes to non-targeting controls as described^47^. Essential and negative regulators of cell growth were identified using an FDR cut-off of 5%.

### RNASeq of WSU-FSCCL IgG^+^ vs IgM^+^ cells

RNA was isolated from 5×10^6^ sorted WSU-FSCCL IgG^+^ and IgM^+^ cells using the RNeasy mini kit (Qiagen, 74004). RNA quality control was performed using a Bioanalyzer (Agilent). Stranded mRNA libraries were prepared a KAPA mRNA HyperPrep Kit (Roche) and sequenced on the Illumina HiSeq 4000 platform (75bp single-end sequencing).

Bulk RNA-seq analysis workflow was performed on LatchBio. First, as a pre-processing step, FastQC was used to confirm per base sequence quality giving the per-site distribution over the length of the read and sequence duplication levels. TrimGalore was used to remove Illumina sequencing adaptors, followed by transcript “pseudo-alignment” using salmon. Tximport was used to convert mapped transcripts into gene-level read counts.

Differential expression analysis was performed with the DESeq2 package. The counts table provided by salmon was used as the input. List of differentially expressed genes and volcano plots were generated with an adjusted p-value of 0.05 as the significance for differentially expressed genes. Pathway analysis on a list of significant differentially expressed genes from DESeq2 analysis was performed using pathfindR^48^.

### Cell surface staining for IgM and IgG cells by flow cytometry

B-cells were washed with FACS buffer consisting of 1x PBS (Gibco), 1% bovine serum albumin (Sigma-Aldrich), and 0.1% sodium azide (Sigma-Aldrich) then immunostained with anti-IgM Fab conjugated with Alexa Fluor 647 (Table 2, Jackson ImmunoResearch), anti-IgG Fab conjugated with DyLight 405 (Jackson ImmunoResearch), anti-Igκ-biotin (Abcam) conjugated in-house with Alexa Fluor 488 NHS Ester (Invitrogen) at a molar ratio of 10:1 dye to antibody, and anti-Igλ conjugated with Alexa Fluor 488 (BioLegend) all at 1:200 v/v for 30 minutes at 4°C (Table 2). Cells were also stained with 1:1,000 dilution of Fixable Viability Dye eFluor™ 780 (eBioscience). Cells were washed and resuspended in FACS buffer for flow analysis using the BD LSRFortessa™. Growth assays were conducted by staining for IgM and IgG in mCherry^+^ transduced cells cultured cells over time.

### Intracellular staining of IgM^+^ and IgG^+^ cells

WSU-FSCCL, NU-DHL1, HBL1, and SUDHL5 IgM^+^ and IgG^+^ were set up in co-culture at equal ratios for at least 24 hours before analysis. Cells were then resuspended in RPMI complete media at 1×10^6^ cells/ml and stimulated with 5 µg/ml of anti-Igκ plus anti-Igλ F(ab’)_2_ antibodies (BioRad, Table 2) for the indicated time at 37°C. Cells were then topped off with cold pFACs buffer (1X HBSS (Gibco), 1% bovine serum albumin (Sigma-Aldrich), 0.1% sodium azide (Sigma-Aldrich)) and stained with at 1:1,000 dilution of Fixable Viability Dye eFluor™ 780 5 min on ice, with all subsequent washing steps performed in pFACs buffer. Cells were then fixed using a 4% paraformaldehyde solution in HBSS (VWR) for 15 mins at RT. Cells were washed again and then resuspended in 0.1% TritonX (Sigma-Aldrich) solution in HBSS for 5 mins at RT. Cells were washed and blocked with 5% BSA in HBSS for 20 mins at RT. Cells were then stained in pFACs buffer for pGSK3, pAKT, pERK, pCD79a, and pCD19 as indicated in Table 2 for 1 hour at RT. For intracellular Ig light chain staining, cells were washed, fixed and permeabilized using the BD Cytofix/Cytoperm™ kit (BD Biosciences). Cells were washed and then stained with anti-IgM Fab and anti-IgG Fab along with anti-rabbit AF488-labeled secondary antibody (Table 2). Cells were washed one final time and resuspended in pFACS buffer for flow cytometry using the BD LSRFortessa™.

### Effect of cell stimulation on growth

WSU-FSCCL and NU-DHL1 IgM^+^ and IgG4^+^ cells were set up in co-culture at equal ratios 24 hours before stimulation. Baseline IgM and IgG cell ratios were analysed by flow cytometry as above. Cultures were then treated with 20 μg/ml DC-SIGN-Fc (R&D Systems), or the indicated concentration of anti-Igκ F(ab’)_2_. Cells were restimulated at day 3 and IgM to IgG cell ratio analysed on day 7.

### Ca^2+^ signalling

WSU-FSCCL, NU-DHL1, HBL1, and SUDHL5 IgM^+^ and IgG^+^ were set up in co-culture at equal ratios for at least 24 hours before analysis. Cells were then loaded with 1 μM Fluo-4, AM (Invitrogen) or 4.5 μM Indo-1, AM (ThermoFisher) at a concentration of 1×10^6^ cells/ml in HBSS with 10% FBS at 37°C for 30 minutes. Cells were washed with HBSS and resuspended in HBSS with 10% FBS and stained using anti-IgM, anti-IgG F_ab_ fragments and a viability dye for 15 mins on ice. Cells were then washed and resuspended in HBSS + 10% FBS. Cells were pre-warmed for 5 mins at 37°C before intracellular Ca^2+^ flux was measured by flow cytometry. After collecting a baseline for 30 s, cells were stimulated with 5 μg/ml anti-Igκ plus anti-Igλ F(ab’)_2_ antibodies (BioRad). After 5 min of data acquisition, 5 μg/ml of Ionomycin (Sigma-Aldrich) was added. Change in Fluo-4 fluorescence intensity was recorded and then plotted using FlowJo (TreeStar). Normalised fold change was determined by dividing fluorescence intensity at each timepoint by the baseline intensity. Indo-1 fluorescence emission was measured using a 355 nm laser and 450/50 (Indo-1 violet) and 530/50 (Indo-1 blue) filter sets on the BD LSRFortessa™. The ratio was calculated as Indo-1 violet/Indo-1 blue.

### GSK3 inhibition

WSU-FSCCL Cas9, HBL1 Cas9, and NU-DHL1 Cas9 cells were transduced with dual-targeting plasmids to switch from IgM to IgG. Baseline IgM and IgG ratios were analysed by flow cytometry at day 3 post-transduction. Cells were then washed and loaded with CFSE. The loaded cells were plated in duplicates (0.5×10^6^ cells/ condition) in complete RPMI with either DMSO or 1 µM GSK3 inhibitor CHIR99021 (Sigma-Aldrich) and cultured for 7 days. IgM and IgG percentages in culture and CFSE dilution were then analysed by flow cytometry, with IgM and IgG percentages normalised to day 3 post-transduction (day 0 treatment).

### Retroviral production and transduction for IgG inducible expression

IgH variable region derived from WSU-FSCCL cells coupled to either the WT or tail-mutant variant W375* of the membrane IgG1 constant region were cloned into the doxycycline-controllable expression vector pRetroX-TetOne™-Puro-EGFP. GP2-293 retroviral packaging cell line were co-transfected with 4.6 μg vector plasmid, 0.54 μg envelope plasmid pMD2.G (Addgene #12259), and 0.99 μg packaging plasmid psPAX2 (Addgene #12260) in Opti-MEM media (ThermoFisher) using TransIT-LT1 transfection reagent (Mirus). Retrovirus was harvested 48 hours post-transfection from cell culture supernatant.

WSU-FSCCL cells were resuspended with harvested retrovirus and 10 µg/ml polybrene (Sigma-Aldrich) then spinfected at 1500 g for 90 min at 4°C. Cells were then selected using 1 μM puromycin for two weeks. Expression of the IgG BCR variants was induced by adding 100 nM doxycycline to the culture.

### Western blotting

Samples were collected and washed twice with PBS. Samples were then resuspended at 1 x 10^6^ cells in 50 μl RIPA buffer (Sigma-Aldrich) with 1X cOmplete™ Protease Inhibitor Cocktail (Roche), incubated for 30 min on ice, before centrifugation for 3 min at 13,000 rpm at 4 °C in a microcentrifuge to separate the cell pellet from cell lysate. 30 μl of cell lysate was mixed with 4X NuPAGE LDS Sample Buffer (Invitrogen) and 10X NuPAGE Sample Reducing Agent (Invitrogen), then boiled for 5 min at 95 °C. Protein samples were loaded onto a NuPAGE 4 - 12% Bis-Tris protein gel (Invitrogen) along with PageRuler Plus Prestained Protein Ladder (Thermo Scientific). The gel was run for 10 min at 100 V then 35 min at 200 V. Protein was transferred onto a PVDF membrane (Bio-Rad) for 35 min at 25 V. Membranes were blocked in 5% skimmed milk in TBST (1X Tris-Buffered Saline, 0.1% Tween 20 Detergent) for 1 hour before incubation with primary antibodies overnight at 4 °C. Membranes were washed thrice with TBST and incubated with secondary antibody (1:10,000 goat anti-rabbit IgG, HRP conjugate) for 1 hour at room temperature. Pierce™ ECL Plus Western Blotting Substrate (Thermo Scientific) was used for signal development according to manufacturer’s instructions. Images were acquired using ChemiDoc (Bio-Rad). Analysis was carried out using Image Lab Software (Bio-Rad) to quantify the background-subtracted protein signal intensity. GAPDH was used as a loading control for normalisation.

## Acknowledgements

We thank the Crick high-throughput sequencing, bioinformatic and flow-cytometry facilities. This work was supported by the Francis Crick Institute, which receives its core funding from Cancer Research UK (CC2006, CC2078), the UK Medical Research Council (CC2006, CC2078) and the Wellcome Trust (CC2006, CC2078), and by a CRUK programme grant (C2750/A23669). S. W. T. C. was supported by an MRC DTP training programme. For the purpose of Open Access, the author has applied a CC BY public copyright licence to any Author Accepted Manuscript version arising from this submission.

## Author contributions

L. W. carried out bioinformatics, designed and executed experiments and co-wrote the manuscript. S. W. T. C. carried out bioinformatics, and designed and executed experiments and co-wrote the manuscript. B. S., L. P. and S. L. carried out bioinformatics. L.Z, H.T. and Z.S.T. carried out mouse xenograft experiments. D.C. designed and supervised mouse experiments. F. F., J. O., and K. T. supervised bioinformatic analyses. N. E. provided reagents and designed experiments. P. T. supervised the study and co-wrote the manuscript. All authors edited the manuscript.

---

**Extended Data Fig. 2.**
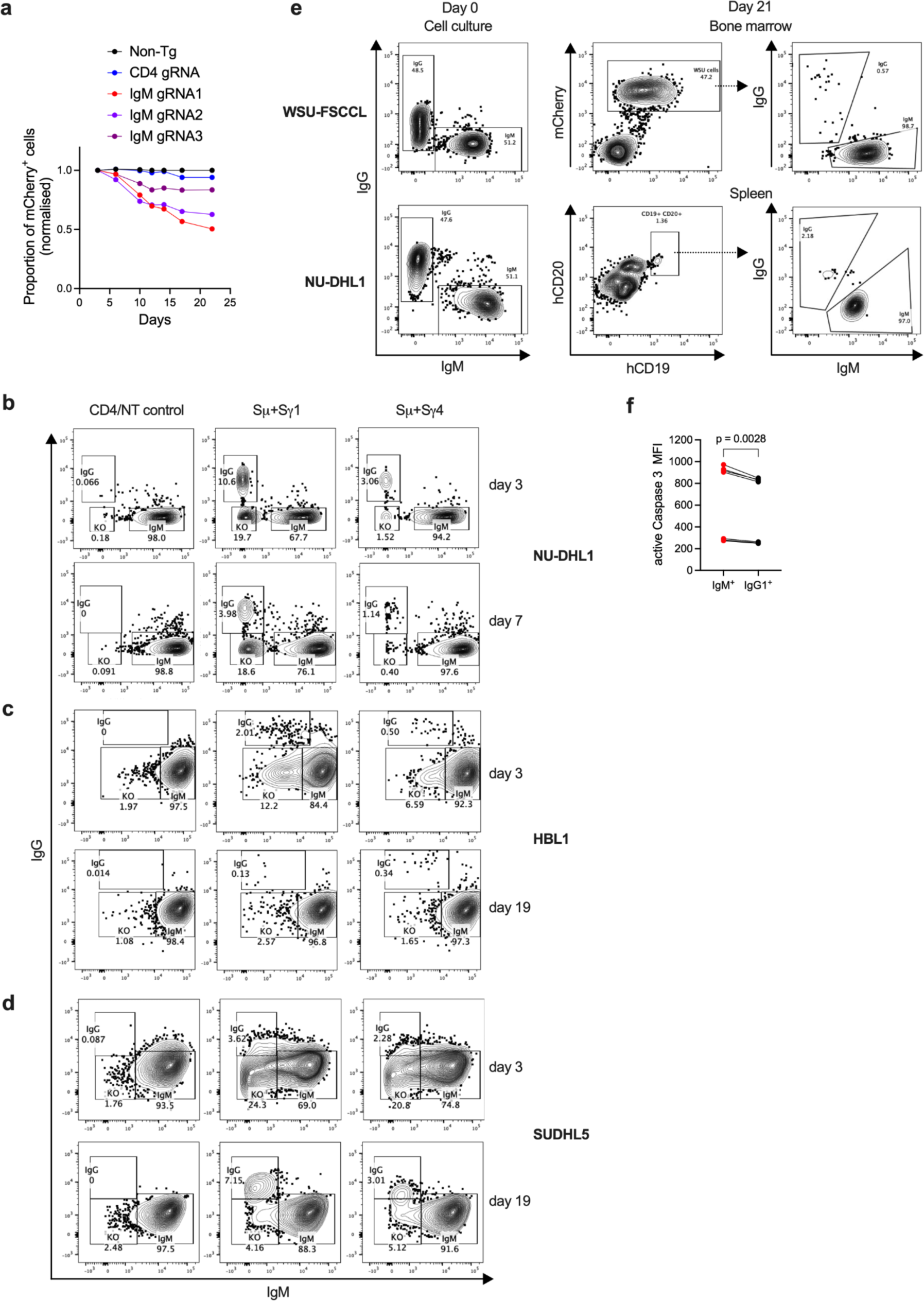
Class-switching from IgM to IgG inhibits the growth of EZB and MCD DLBCL cell lines. (a) Proportions of WSU-FSCCL Cas9 cells targeted with the indicated sgRNA-containing lentiviruses expressing mCherry. One experiment out of three biological replicates is shown. (b-d) Representative flow cytometry plots showing staining with anti-IgM and anti-IgG BCR in the indicated cell lines after Cas9/sgRNA nucleofection (b) or lentiviral delivery (c, d) of dual sgRNAs targeting the indicated genomic regions. Proportions of IgM^+^, IgM^-^ IgG^-^ and IgG^+^ cells are shown on day 3 and day 7 or 19 after class-switching. (e) Representative flow cytometry detecting IgG1^+^ and IgM^+^ WSU-FSCCL or IgG1^+^ and IgM^+^ NU-DHL1 cells mixed 1:1 before (left) and 3 weeks after xenotransplantation into NSG mice (right). Gates used to detect transplanted cells in the bone marrow (WSU-FSCCL) or spleen (NU-DHL1) are shown in the middle panels. (f) Proportions of Caspase 3^+^ cells in sorted IgG4^+^ and IgM^+^ WSU-FSCCL cells. Bracket shows significance in a paired t-test. N = 8 experiments.

**Extended Data Fig. 3.**
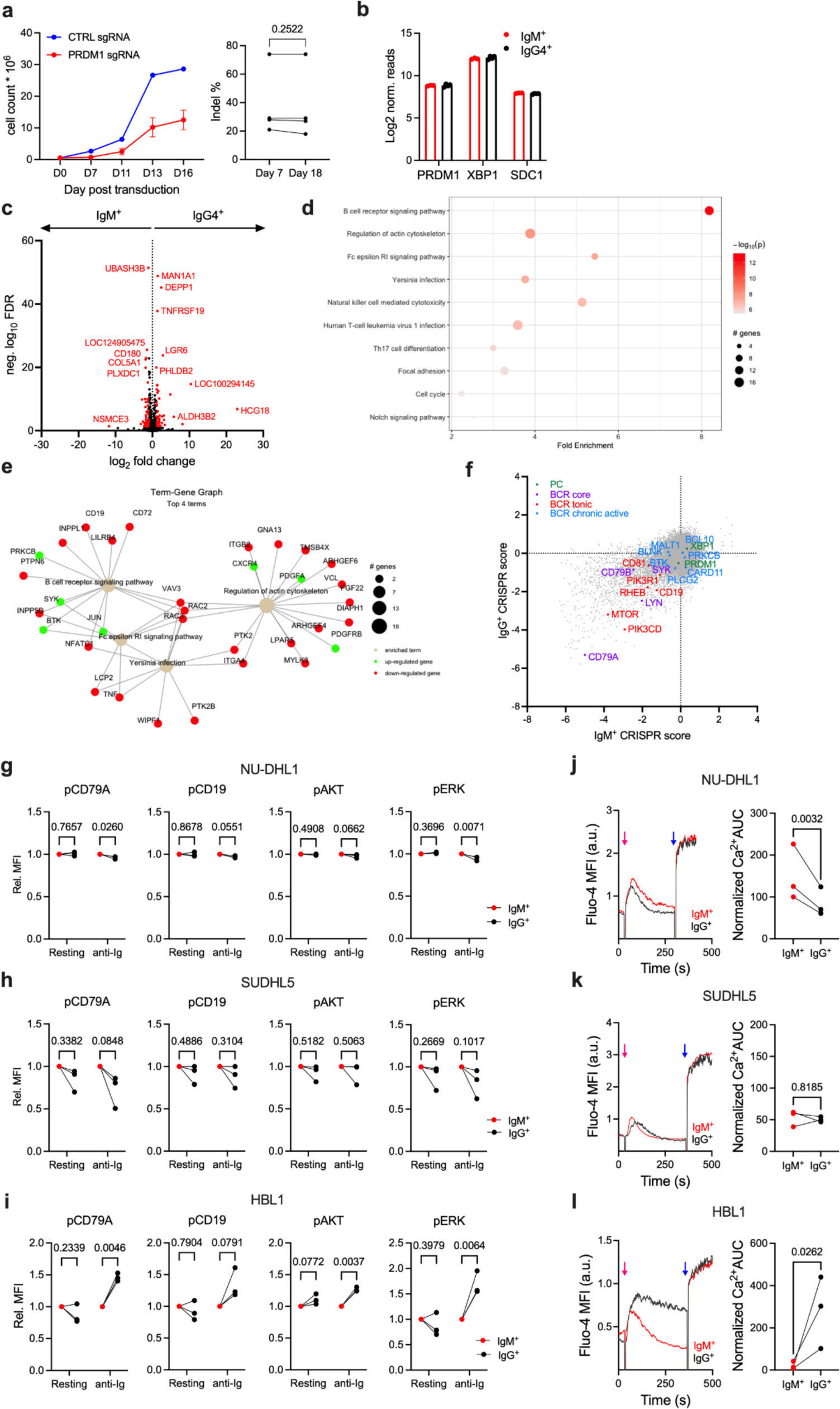
Class-switching modulates lymphoma BCR signalling but not PC differentiation. (a) Targeting PRDM1 in IgG4^+^ WSU-FSCCL does not rescue cell growth. Left, growth curves of cells transduced with non-targeting or PRDM1-targeting sgRNAs along with puromycin resistance in puromycin-containing media. Right, the frequency of indels at the PRDM1 locus does not show an accumulation of PRDM1-targeted cells in the culture. Brackets show significance in a paired t-test, N = 4 experiments. (b-d) RNAseq comparison of IgM^+^ and IgG4^+^ WSU-FSCCL cells. N = 4 experiments. (b) Expression of PC differentiation genes PRDM1 and XB1. (c) Volcano plot comparing transcripts in IgM^+^ and IgG4^+^ cells. (d) Pathway enrichment in gene expression differences. (e) Gene expression difference in the four top differentially expressed pathways. (f) CRISPR scores determined in whole-genome CRISPR screens for genes regulating the growth of IgM^+^ and IgG4^+^ WSU-FSCCL cells. N = 2 replicates. Gene sets associated with BCR signalling and PC differentiation are highlighted. (g-i) Comparison of the levels of phopho-CD79A, phospho-CD19, phospho-AKT and phospho-ERK in IgM^+^ and IgG4^+^ cells of the indicated cell lines before and 10 min after stimulation with anti-Ig light chain antibodies as determined by intracellular staining and flow cytometry. Data show the mean fluorescence intensity of cells in each experiment normalised to values of the IgM^+^ cells. Brackets show P values in repeat-measures two-way ANOVA, N = 3 experiments. (j-l) Left, intracellular Ca^2+^ levels in IgM^+^ and IgG4^+^ cells as determined by Fluo-4 fluorescence after stimulation with anti-Ig (red arrow) and ionomycin (blue arrow). Right, quantification of the area under the Ca^2+^ curve covering anti-Ig stimulation. P values in paired t-test, N = 3 experiments.

**Extended Data Fig. 4.**
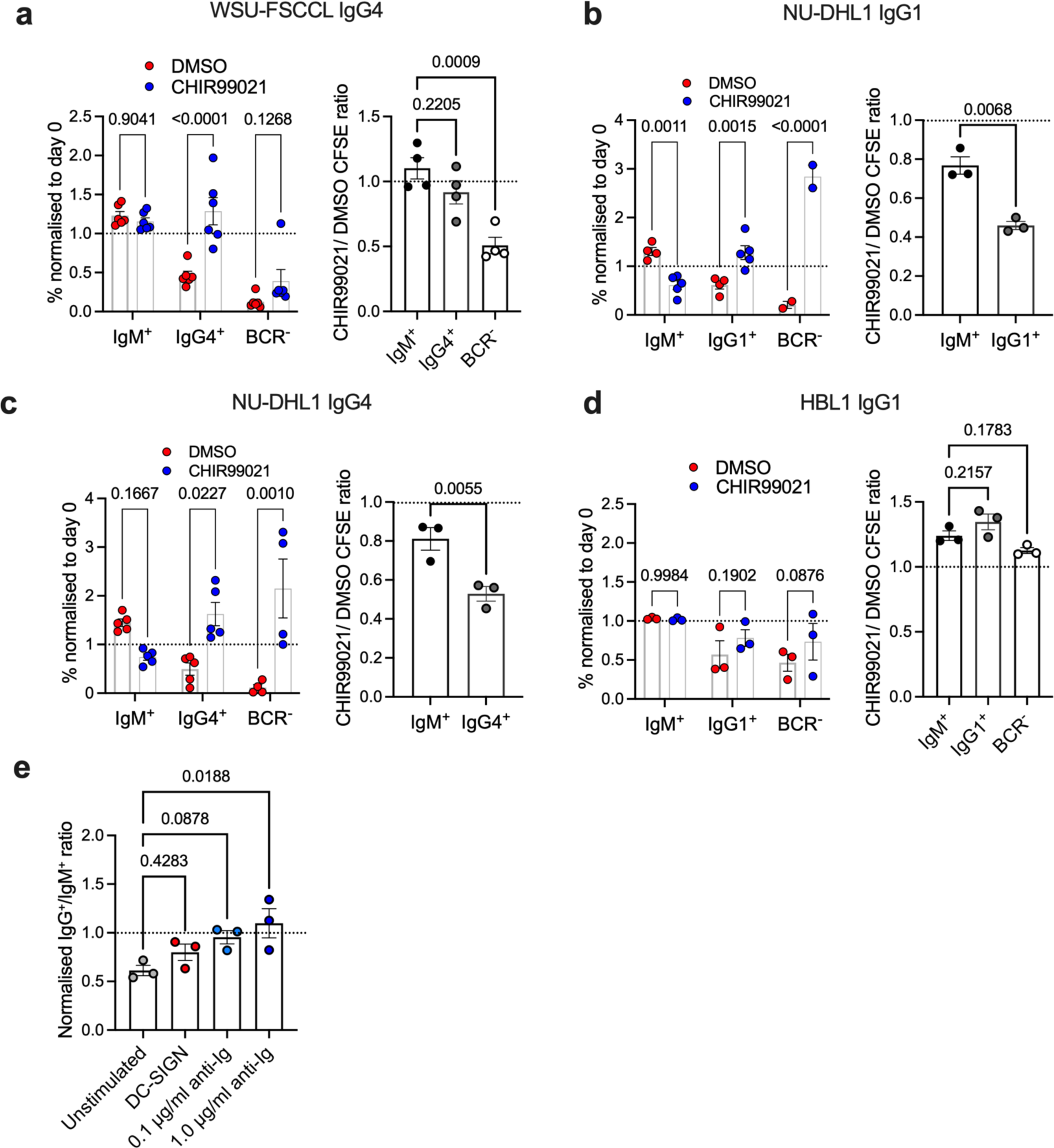
Bypassing BCR signalling to GSK3 or BCR stimulation rescues IgG-switched EZB DLBCL cells. (a-d) Left panels show the percent (normalised to day 0) of IgM^+^, IgG1^+^ and BCR^-^ WSU-FSCCL cells 7 days after CRISPR-mediated class-switching in the presence of vehicle (DMSO) or GSK3 inhibitor (CHIR99021). Right panels show the effect on CFSE dilution. Brackets show P values in two-way ANOVA, N = 2 to 6 experiments. (e) Growth of IgM^+^ and IgG4^+^ NU-DHL1 cells after stimulation. Data show the ratio of IgG^+^ to IgM^+^ cells after 7 days of stimulation, normalised to day 0. Brackets show P values in two-way ANOVA pairwise comparisons to unstimulated cells, N = 3 experiments.

**Extended Data Fig. 5.**
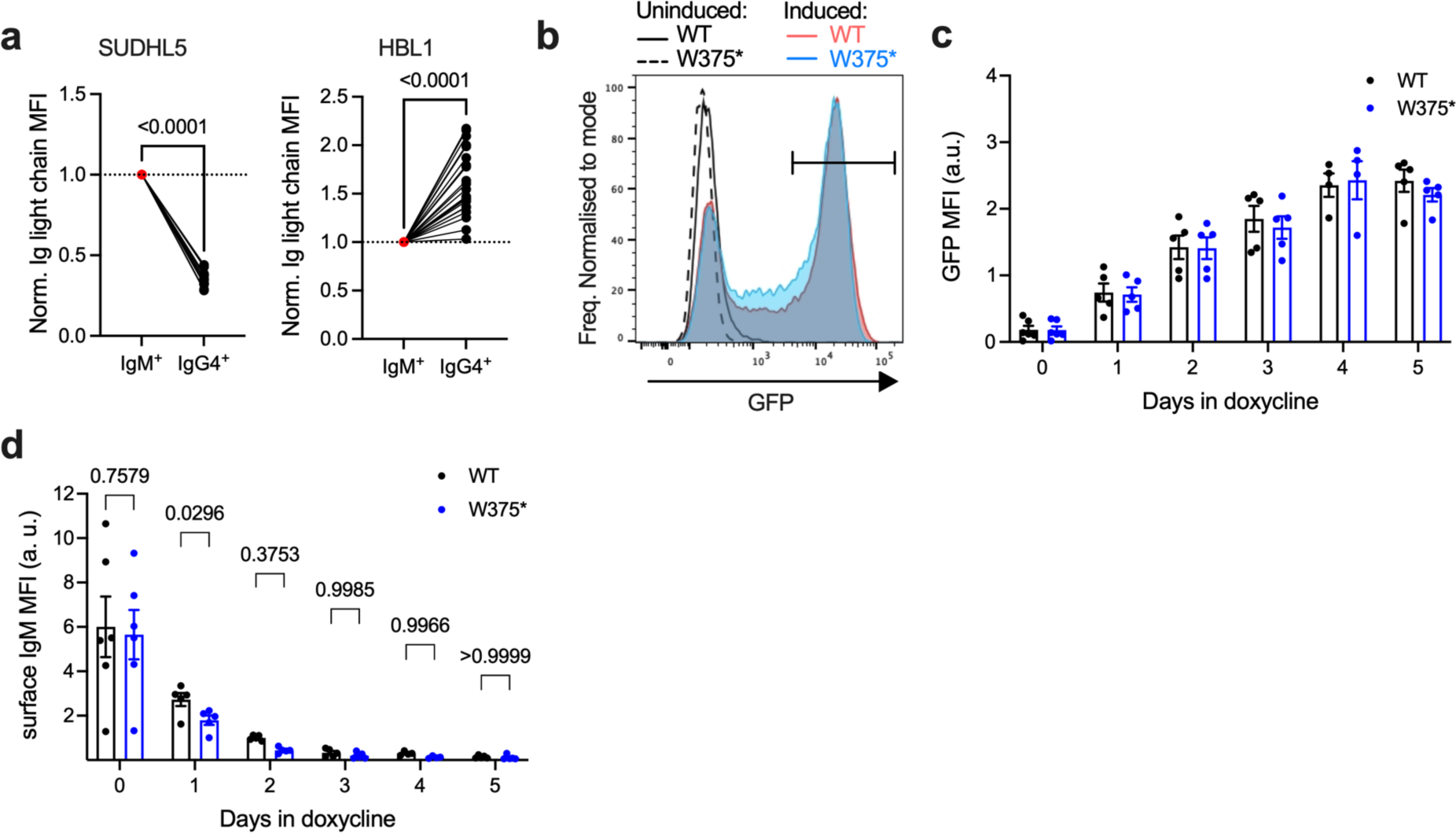
IgG tail regulates BCR surface levels and signalling. (a) Surface BCR levels in IgM^+^ and IgG4^+^ SUHL5 and HBL1 cells, normalised to IgM^+^ levels, as determined by anti-Ig light chain staining and flow cytometry. Brackets show P values in paired t-tests, N = 5 or 25. (b) Representative flow cytometry histograms of GFP levels in WSU-FSCCL cells transduced with constructs expressing WT or W375* mutated IgG1 containing the endogenous variable domain and bicistronic GFP. Grey and dotted histograms show cells without doxycycline-induced expression, blue and red histograms show cells 2 days after doxycycline-induced expression. (c) GFP MFI of GFP^+^ cells after culture with doxycycline for the indicated number of days. Data show means and SEM, N = 4 to 6 experiments. (d) Surface IgM levels in GFP^+^ cells after culture with doxycycline for the indicated number of days. Data show means and SEM, N = 4 to 6 experiments. Brackets show significance of comparisons in two-way ANOVA.

## Supplementary Data

**Supplementary Table 1.** List of all IgG tail mutations identified in all datasets.

**Supplementary Table 2.** Log2 normalised read counts from RNAseq of IgM^+^ and IgG4^+^ WSU-FSCCL cells.

**Supplementary Table 3.** Results of gene-essentiality CRISPR screens of IgM^+^ and IgG4^+^ WSU-FSCCL cells. CRISPR scores are shown as log2 fold changes (lfc).

## References

1. Lutz, J. et al. Reactivation of IgG-switched memory B cells by BCR-intrinsic signal amplification promotes IgG antibody production. Nat Commun 6, 8575 (2016).

2. Engels, N. et al. Recruitment of the cytoplasmic adaptor Grb2 to surface IgG and IgE provides antigen receptor–intrinsic costimulation to class-switched B cells. Nat Immunol 10, 1018–1025 (2009).

3. Sundling, C. et al. Positive selection of IgG+ over IgM+ B cells in the germinal center reaction. Immunity 54, 988–1001.e5 (2021).

4. Martin, S. W. & Goodnow, C. C. Burst-enhancing role of the IgG membrane tail as a molecular determinant of memory. Nat Immunol 3, 182–188 (2002).

5. Shaffer, A. L., Young, R. M. & Staudt, L. M. Pathogenesis of Human B *Cell* Lymphomas. Annu Rev Immuno. 30, 565–610 (2012).

6. Roulland, S. et al. Chapter 1 Early Steps of Follicular Lymphoma Pathogenesis. Adv Immunol 111, 1–46 (2011).

7. Chapuy, B. et al. Molecular subtypes of diffuse large B cell lymphoma are associated with distinct pathogenic mechanisms and outcomes. Nat Med 24, 679–690 (2018).

8. Wright, G. W. et al. A Probabilistic Classification Tool for Genetic Subtypes of Diffuse Large B Cell Lymphoma with Therapeutic Implications. Cancer Cell 37, 551–568.e14 (2020).

9. Lacy, S. E. et al. Targeted sequencing in DLBCL, molecular subtypes, and outcomes: a Haematological Malignancy Research Network report. Blood 135, 1759–1771 (2020).

10. Crouch, S. et al. Molecular subclusters of follicular lymphoma: a report from the UK’s Haematological Malignancy Research Network. Blood Adv 6, 5716–5731 (2022).

11. Sundling, C. et al. Positive selection of IgG+ over IgM+ B cells in the germinal center reaction. Immunity 54, 988–1001.e5 (2021).

12. Schmitz, R. et al. Genetics and Pathogenesis of Diffuse Large B-Cell Lymphoma. N Engl J Med 378, 1396–1407 (2018).

13. Young, R. M. et al. Taming the Heterogeneity of Aggressive Lymphomas for Precision Therapy. Annu Rev Cancer Biology 3, 1–27 (2018).

14. Vaandrager, J.-W. et al. DNA Fiber Fluorescence In Situ Hybridization Analysis of Immunoglobulin Class Switching in B-Cell Neoplasia: Aberrant CH Gene Rearrangements in Follicle Center-Cell Lymphoma. Blood 92, 2871–2878 (1998).

15. Cha, S.-C. et al. Nonstereotyped Lymphoma B Cell Receptors Recognize Vimentin as a Shared Autoantigen. J Immunol 190, 4887–4898 (2013).

16. Ruminy, P. et al. The isotype of the BCR as a surrogate for the GCB and ABC molecular subtypes in diffuse large B-cell lymphoma. Leukemia 25, 681–688 (2011).

17. Lenz, G. et al. Aberrant immunoglobulin class switch recombination and switch translocations in activated B cell–like diffuse large B cell lymphoma. J Exp Med 204, 633– 643 (2007).

18. Kaisho, T., Schwenk, F. & Rajewsky, K. The Roles of γ1 Heavy Chain Membrane Expression and Cytoplasmic Tail in IgG1 Responses. Science 276, 412–415 (1997).

19. Engels, N. et al. The immunoglobulin tail tyrosine motif upgrades memory-type BCRs by incorporating a Grb2-Btk signalling module. Nat Commun 5, 5456 (2014).

20. Vanshylla, K. et al. Grb2 and GRAP connect the B cell antigen receptor to Erk MAP kinase activation in human B cells. Sci Rep 8, 4244 (2018).

21. Chen, X. et al. An autoimmune disease variant of IgG1 modulates B cell activation and differentiation. Science 362, 700–705 (2018).

22. Reddy, A. et al. Genetic and Functional Drivers of Diffuse Large B Cell Lymphoma. Cell 171, 481–494.e15 (2017).

23. Hübschmann, D. et al. Mutational mechanisms shaping the coding and noncoding genome of germinal center derived B-cell lymphomas. Leukemia 35, 2002–2016 (2021).

24. Klijn, C. et al. A comprehensive transcriptional portrait of human cancer cell lines. Nat Biotechnol 33, 306–312 (2015).

25. Chiodin, G. et al. Insertion of atypical glycans into the tumor antigen-binding site identifies DLBCLs with distinct origin and behavior. Blood 138, 1570–1582 (2021).

26. Salles, G. et al. Rituximab maintenance for 2 years in patients with high tumour burden follicular lymphoma responding to rituximab plus chemotherapy (PRIMA): a phase 3, randomised controlled trial. Lancet 377, 42–51 (2011).

27. Cheong, T.-C., Compagno, M. & Chiarle, R. Editing of mouse and human immunoglobulin genes by CRISPR-Cas9 system. Nat Commun 7, 10934 (2016).

28. Zhang, L. et al. Regulation of BCR-mediated Ca2+ mobilization by MIZ1-TMBIM4 safeguards IgG1+ GC B cell–positive selection. Sci. Immunol. 9, eadk0092 (2024).

29. Havranek, O. et al. Tonic B-cell receptor signaling in diffuse large B-cell lymphoma. Blood 130, 995–1006 (2017).

30. Varano, G. et al. The B-cell receptor controls fitness of MYC-driven lymphoma cells via GSK3β inhibition. Nature 546, 302–306 (2017).

31. McCann, K. J. et al. Remarkable selective glycosylation of the immunoglobulin variable region in follicular lymphoma. Mol Immunol 45, 1567–1572 (2008).

32. Linley, A. et al. Lectin binding to surface Ig variable regions provides a universal persistent activating signal for follicular lymphoma cells. Blood 126, 1902–1910 (2015).

33. Coelho, V. et al. Glycosylation of surface Ig creates a functional bridge between human follicular lymphoma and microenvironmental lectins. Proc Natl Acad Sci 107, 18587–18592 (2010).

34. Pape, K. A., Taylor, J. J., Maul, R. W., Gearhart, P. J. & Jenkins, M. K. Different B Cell Populations Mediate Early and Late Memory During an Endogenous Immune Response. Science 331, 1203–1207 (2011).

35. Dogan, I. et al. Multiple layers of B cell memory with different effector functions. Nat Immunol 10, 1292–1299 (2009).

36. Zuccarino-Catania, G. V. et al. CD80 and PD-L2 define functionally distinct memory B cell subsets that are independent of antibody isotype. Nat Immunol 15, 631–637 (2014).

37. Young, R. M. et al. Survival of human lymphoma cells requires B-cell receptor engagement by self-antigens. Proc National Acad Sci 112, 13447–13454 (2015).

38. Kodama, T., Hasegawa, M., Sakamoto, Y., Haniuda, K. & Kitamura, D. Ubiquitination of IgG1 cytoplasmic tail modulates B-cell signalling and activation. Int Immunol 32, 385–395 (2020).

39. Amin, R. et al. DC-SIGN–expressing macrophages trigger activation of mannosylated IgM B-cell receptor in follicular lymphoma. Blood 126, 1911–1920 (2015).

40. Horikawa, K. et al. Enhancement and suppression of signaling by the conserved tail of IgG memory-type B cell antigen receptors. J Exp Med 204, 759–769 (2007).

41. Gitlin, A. D. et al. Independent Roles of Switching and Hypermutation in the Development and Persistence of B Lymphocyte Memory. Immunity 44, 769–781 (2016).

42. Davis, R. E. et al. Chronic active B-cell-receptor signalling in diffuse large B-cell lymphoma. Nature 463, 88–92 (2010).

43. Preece, R. et al. ‘Mini’ U6 Pol III promoter exhibits nucleosome redundancy and supports multiplexed coupling of CRISPR/Cas9 effects. Gene Ther. 27, 451–458 (2020).

44. Shy, B. R. et al. High-yield genome engineering in primary cells using a hybrid ssDNA repair template and small-molecule cocktails. Nat Biotechnol 41, 521–531 (2023).

45. Clement, K. et al. CRISPResso2 provides accurate and rapid genome editing sequence analysis. Nat Biotechnol 37, 224–226 (2019).

46. Li, W. et al. Quality control, modeling, and visualization of CRISPR screens with MAGeCK-VISPR. Genome biology 16, 281 (2015).

47. Newman, R. & Tolar, P. Chronic calcium signaling in IgE+ B cells limits plasma cell differentiation and survival. Immunity 54, 2756–2771.e10 (2021).

48. Ulgen, E., Ozisik, O. & Sezerman, O. U. pathfindR: An R Package for Comprehensive Identification of Enriched Pathways in Omics Data Through Active Subnetworks. Front Genet 10, 858 (2019).

